# Redox sensitive E2 Rad6 controls cellular response to oxidative stress via K63 ubiquitination of ribosomes

**DOI:** 10.1101/2021.04.22.439970

**Authors:** Vanessa Simões, Lana Harley, Blanche K. Cizubu, Ye Zhou, Joshua Pajak, Nathan A. Snyder, Jonathan Bouvette, Mario J. Borgnia, Gaurav Arya, Alberto Bartesaghi, Gustavo M. Silva

## Abstract

Protein ubiquitination is an essential process that rapidly regulates protein synthesis, function, and fate in dynamic environments. Among its non-proteolytic functions, K63 ubiquitin accumulates in yeast cells exposed to oxidative stress, stalling ribosomes at elongation. K63 ubiquitin conjugates accumulate because of redox inhibition of the deubiquitinating enzyme Ubp2, however, the role and regulation of ubiquitin conjugating enzymes in this pathway remained unclear. Here we found that the E2 Rad6 binds and modifies elongating ribosomes during oxidative stress. We elucidated a mechanism by which Rad6 and its human homolog UBE2A are redox-regulated by forming reversible disulfides with the E1 activating enzyme, Uba1. We further showed that Rad6 activity is necessary to regulate translation, antioxidant defense, and adaptation to stress. Finally, we showed that Rad6 is required to induce phosphorylation of the translation initiation factor eIF2α, providing a novel link for K63 ubiquitin, elongation stalling, and the integrated stress response.

## Introduction

Translation is a highly regulated process in eukaryotes, controlled at multiple steps during cellular exposure to stress^1,2^. Using mass spectrometry and cryo-electron microscopy (cryo-EM), our group recently showed that oxidative stress induces high levels of K63 ubiquitinated ribosomes, which impact the elongation step of translation^3–5^. We named this pathway RTU: Redox control of Translation by Ubiquitin^6^. Protein ubiquitination is a multi-step process and much remains unknown about the regulation of K63 ubiquitination of ribosomes in the RTU. In general, protein ubiquitination is achieved by the activity of ubiquitin conjugases (E2s) and ligases (E3s), which can be counteracted by deubiquitinases (DUBs) and degradation processes^7^. As the majority of ubiquitin enzymes contain cysteine residues in their catalytic sites, these enzymes can undergo redox modifications and have their activity regulated by reactive oxygen species (ROS)^8,9^. In the RTU, we determined that redox inhibition of the DUB Ubp2 is essential for the accumulation of K63 ubiquitin conjugates under stress^3^. However, how ubiquitin conjugating enzymes contribute to K63 ubiquitination of ribosomes and how their activity is regulated in response to oxidative stress still remain unknown.

In the canonical ubiquitination pathway, the E2-E3 interaction determines substrate specificity^10^, while the E2 is largely responsible for the ubiquitin chain topology extended on these substrates^11^. We have previously shown that deletion of the E2 *RAD6* was the sole genetic modification that prevented the accumulation of K63 ubiquitin after cellular exposure to hydrogen peroxide (H_2_O_2_)^3,4^. Interestingly, it has been previously suggested that Rad6 would be able to transfer K63 ubiquitin chains *en bloc* to its substrate PCNA (Proliferating Cell Nuclear Antigen), however, these K63 ubiquitin chains were built by Ubc13^12^, the main K63 ubiquitin E2 in yeast^13^. Deletion of Ubc13 did not prevent K63 ubiquitin accumulation under stress^3^, which suggests that additional E2’s would be able to conjugate K63 ubiquitin chains to ribosomes during stress. Rad6 is a highly conserved, multifunctional E2^14^, in which several mutations to its human homolog (UBE2A) have been linked to learning and cognitive disabilities^15,16^. Moreover, Rad6 has been described to modify targets with mono-ubiquitin or K48 ubiquitin, being involved in DNA repair, the N-end rule pathway, and transcriptional control^17–19^, interacting in each of these pathways with the E3’s Rad18, Ubr1, and Bre1, respectively. We have shown that deletion of *BRE1* also hindered the accumulation of K63 ubiquitin conjugates in response to stress^3^, suggesting a new function for this E2-E3 pair. However, it remained unclear whether Rad6 directly modifies ribosomes and whether its activity is regulated by ROS upon exposure to oxidative stress. Furthermore, we lacked understanding on how deletion of *RAD6* impacts translation and cellular resistance to stress.

In this study, we set out to investigate the key role of Rad6 in regulating cellular response to stress as part of the RTU. First, we found that Rad6 promotes K63 ubiquitination of ribosomes *in vitro* and *in vivo*. We also determined a mechanism by which, following ubiquitin conjugation, Rad6 is redox regulated by forming a reversible disulfide with the E1 activating enzyme, Uba1. We further showed that Rad6 function in the K63 ubiquitination of ribosomes can be complemented by UBE2A, which can also be redox-regulated by disulfide formation in human cells. Finally, we showed that Rad6 is necessary for reprogramming translation and for adequate synthesis of antioxidant proteins, suggesting a new mechanism for the activation of the integrated stress response.

## Results

### Rad6 is the E2 responsible for ribosomal K63 ubiquitination

By examining the roles of ubiquitin conjugating enzymes in the RTU, we determined that deletion of *RAD6* in budding yeast diminished the levels of K63 ubiquitinated ribosomes that occur in response to oxidative stress induced by H_2_O_2_ (Supplementary Fig. 1a,b). To further understand the role of Rad6 in K63 ubiquitination of ribosomes, we tested Rad6’s ability to directly bind and ubiquitinate ribosomes. We first showed that the expression of *RAD6* in the *rad6Δ* mutant strain is sufficient to recover both its growth defect and the ribosomal K63 ubiquitin levels (Fig. 1a, b and Supplementary Fig. 1c, d). Next, we reconstituted an *in vitro* ubiquitination system, which showed that affinity-purified Rad6 is able to directly polyubiquitinate ribosomes (Fig. 1c and Supplementary Fig. 2a). Rad6-mediated ubiquitination of ribosomes *in vitro* was also observed in the absence of Bre1 (Fig. 1c), however, deletion of Bre1 drastically reduces the levels of K63 ubiquitinated ribosomes in the cellular context (Supplementary Fig. 2b). Ubiquitination reactions were performed with ribosomes isolated from a *rad6*Δ*bre1*Δ strain, and were also controlled for Rad6 auto-ubiquitination and for residual activity of Ubc13 and Hel2 (Supplementary Fig. 2c-f). Ubc13 and Hel2 are known ubiquitin enzymes involved in K63 ubiquitination and ubiquitination of ribosomes in quality control pathways, respectively^13,20^. We also confirmed that Rad6 can be charged and transfer K63-linked di-ubiquitin to ribosomes (Supplementary Fig. 2g-h). Thus, our results provide direct evidence for a new role of Rad6 in ubiquitinating ribosomes and also reveal a new E2 enzyme able to participate in K63 ubiquitin conjugation reactions.

**Figure 1.**
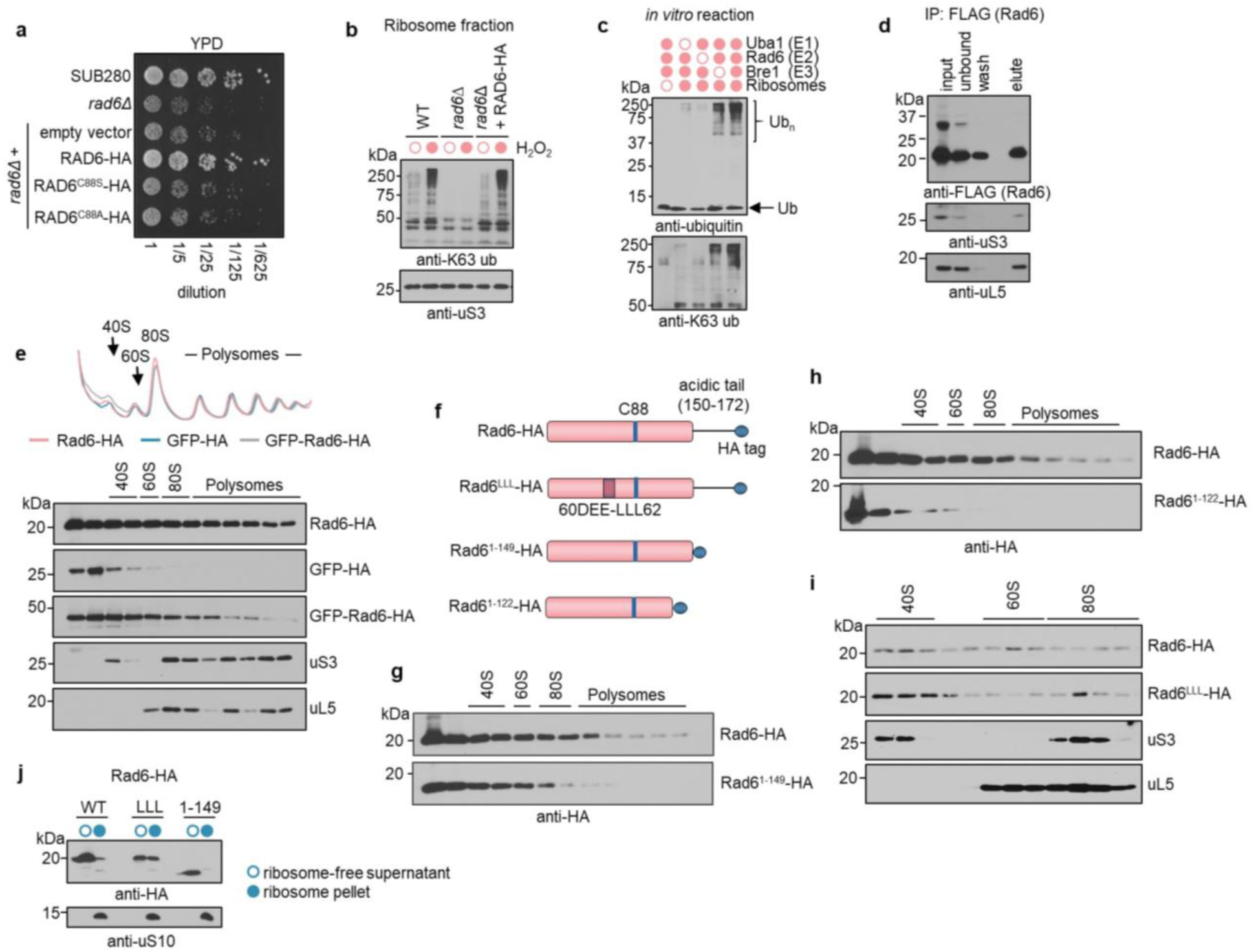
Rad6 binds and ubiquitinates ribosomes. **a**,**b**, Expression of Rad6-HA reverts *rad6Δ* growth defect (**a**) and supports K63 ubiquitin accumulation under 0.6 mM H_2_O_2_ visualized by immunoblotting isolated ribosomes (**b**). **c**, Immunoblot from *in vitro* ubiquitination assay. **d**, Immunoblot of co-immunoprecipitation (IP) of ribosomal protein uS3 and uL5 with Rad6-FLAG. **e**, Sucrose sedimentation profile and immunoblot of lysates from cells expressing Rad6-HA, GFP-HA, and GFP-Rad6-HA constructs. uS3 and uL5 were used as markers for the 40S and 60S ribosome subunits, respectively. **f**, Schematic of Rad6 constructs. **g**,**h**, anti-HA (Rad6) immunoblot from sucrose sedimentation profile shows differential ribosomal binding pattern for Rad6^1-149^ (**g**), Rad6^1-122^ (**h**), and Rad6^LLL^ variants (**i**). **j**, Immunoblot from supernatant (open circle) and ribosome pellet (full circle) after sucrose cushion from strains expressing Rad6^1-149^ and Rad6^LLL^ variants. uS10 was used as ribosome marker.

We next sought to provide insights into the mechanisms by which Rad6 interacts with ribosomes to promote ubiquitination in a cellular context. Our previous data showed that fully assembled ribosomes can be ubiquitinated in response to stress^5^. We then confirmed that Rad6 binds to ribosomes by immunoprecipitating FLAG-tagged Rad6 and detecting the presence of proteins from the small 40S and large 60S subunits of the ribosome (Fig. 1d). By performing polysome profiling analysis, we identified Rad6 present in the light (40S, 60S, 80S) and heavy fractions (polysome) of the sucrose gradient (Fig. 1e). This co-sedimentation pattern is independent of the presence or location of the HA-tag (Supplementary Fig. 3a-b). Rad6 incubated with isolated ribosomes or fused to GFP, relocated these proteins to polysomes (Fig. 1e, Supplementary Fig. 3c). Rad6 interaction with ribosomes is strong and is not impacted by high salt concentration, thiol reducing agents, DNAse, RNAse, or DUB treatment. We also showed that Rad6 binding to ribosomes is not exclusively dependent on Rad6’s known E3’s (Bre1, Ubr1, and Rad18) (Supplementary Fig. 3d-g), further indicating that Rad6 binds directly to ribosomes and not RNA, ubiquitin, or other associated proteins. Finally, we observed that Bre1, the RTU E3 ligase^3^, is also present in the polysome fraction (Supplementary Fig. 3h), supporting a model that the entire enzymatic system is associated with ribosomes to promote K63 ubiquitination.

The presence of Rad6 in the heavier fractions of the profile of unstressed cells suggests that Rad6 interacts constitutively with ribosomes instead of being actively recruited during oxidative stress. Rad6 recruitment to polysomes is independent of stress induction and of Rad6’s catalytic cysteine (C88) (Supplementary Fig. 3i-j). By targeting many of Rad6’s known domains and important residues^21^, we showed that Rad6 binding to ribosomes is not affected by mutations that disrupt its canonical (S25) or the backside ubiquitin binding domain (L56) (Supplementary Fig. 3k). However, removal of Rad6 acidic tail (denoted as Rad6^1–149^), which is suggested to stabilize its interaction with substrates^22,23^, decreased Rad6 binding to polysomes (Fig. 1g). Subsequent removal of Rad6 last alpha helix (denoted as Rad6^1-122^) significantly reduced its presence in the heavy polysome fraction (Fig. 1h). Moreover, mutation to Rad6 acidic region (DEE60-62LLL, hereafter named Rad6^LLL^), which has been proposed to impact the E2-E3 interaction surface^24^, fostered Rad6 association with ribosomes, with increased localization to the 80S (Fig. 1i). Both Rad6^1-149^ and Rad6^LLL^ are still capable of ubiquitinating ribosomes and complement the growth sensitivity of *rad6Δ* (Supplementary Fig. 4a-d). However, deletion of Rad6’s last alpha helix (Rad6^1-122^) prevented K63 ubiquitination and impacted cellular growth, suggesting an important role of the enzyme C-terminus on ribosome binding and function during stress (Supplementary Fig. 4e-g).

Next, we showed that Rad6 is able to bind 80S ribosomes at the pre-translocation stage of elongation. Single-particle cryo-EM analysis of isolated ribosomes incubated *in vitro* with purified Rad6 (Ribo^Rad6mix^) showed five distinct classes distributed along different stages of elongation (Fig. 2a, Supplementary Fig. 5). In this system, we were not able to observe density for Rad6, likely because of its small size and intrinsic flexibility. To gain further understanding from a physiological standpoint, we co-immunoprecipitated ribosomes natively bound to FLAG-tagged Rad6 (Ribo^Rad6IP^) and subjected this sample to cryo-EM imaging. Our analysis revealed that Rad6-bound ribosomes were present at the classic and the rotated pre-translocation stage of translation, containing densities for two tRNAs in the hybrid A/P and P/E positions (Fig. 2a,b). Using focused classification, we identified two sub-classes for the 40S beak with a conventional and extended version of the 18S rRNA, displaying a reorganization of the proteins eS31 and eS12 (Fig. 2c). Focused classification of Ribo^Rad6mix^ showed a dynamic motion of the 40S beak including the K6 class with its extended beak (Supplementary Fig. 6). To increase stability of the complex, we cross-linked the immunoprecipitated ribosomes bound to Rad6 (Ribo^Rad6IP-XL^), which also revealed the extended form of the 40S beak (Supplementary Fig. 6). We also compared Ribo^Rad6IP^ with the structure of K63 ubiquitinated ribosomes and ribosomes from the K63R ubiquitin mutant strain^5^. Although we saw the same extended 18S rRNA conformation in all the 3 datasets (Supplementary Fig.6 d, e), we observed a more dynamic behavior of the 40S beak (elongated density and multiple conformations) on K63 ubiquitinated ribosomes while the K63R mutant dataset showed a combination of conventional and extended 40S beak conformations (Supplementary 6d,e). To gain further insights on Rad6 interaction with ribosomes, we carried out molecular docking calculations and inspected the resulting models on the 40S beak, which showed distinctive conformational changes by cryo-EM and is highly ubiquitinated under stress^4^. By considering constraints, such as the requirement for the Rad6 catalytic cysteine to be proximal to the ubiquitinated lysine, we narrowed down the top scoring models to a single, viable docking pose for each of the ubiquitin sites on the ribosomal proteins (Fig. 2d,e and Supplementary Fig. 7). Interestingly, in most cases the candidate poses placed the acidic 60-DEE-62 region of Rad6 close to hydrophobic patches on the ribosome surface and predicted interactions between the C-terminal residues of Rad6 and the ribosome, providing further support for our experimental findings with mutants Rad6^LLL^ and Rad6^1-122^.

**Figure 2.**
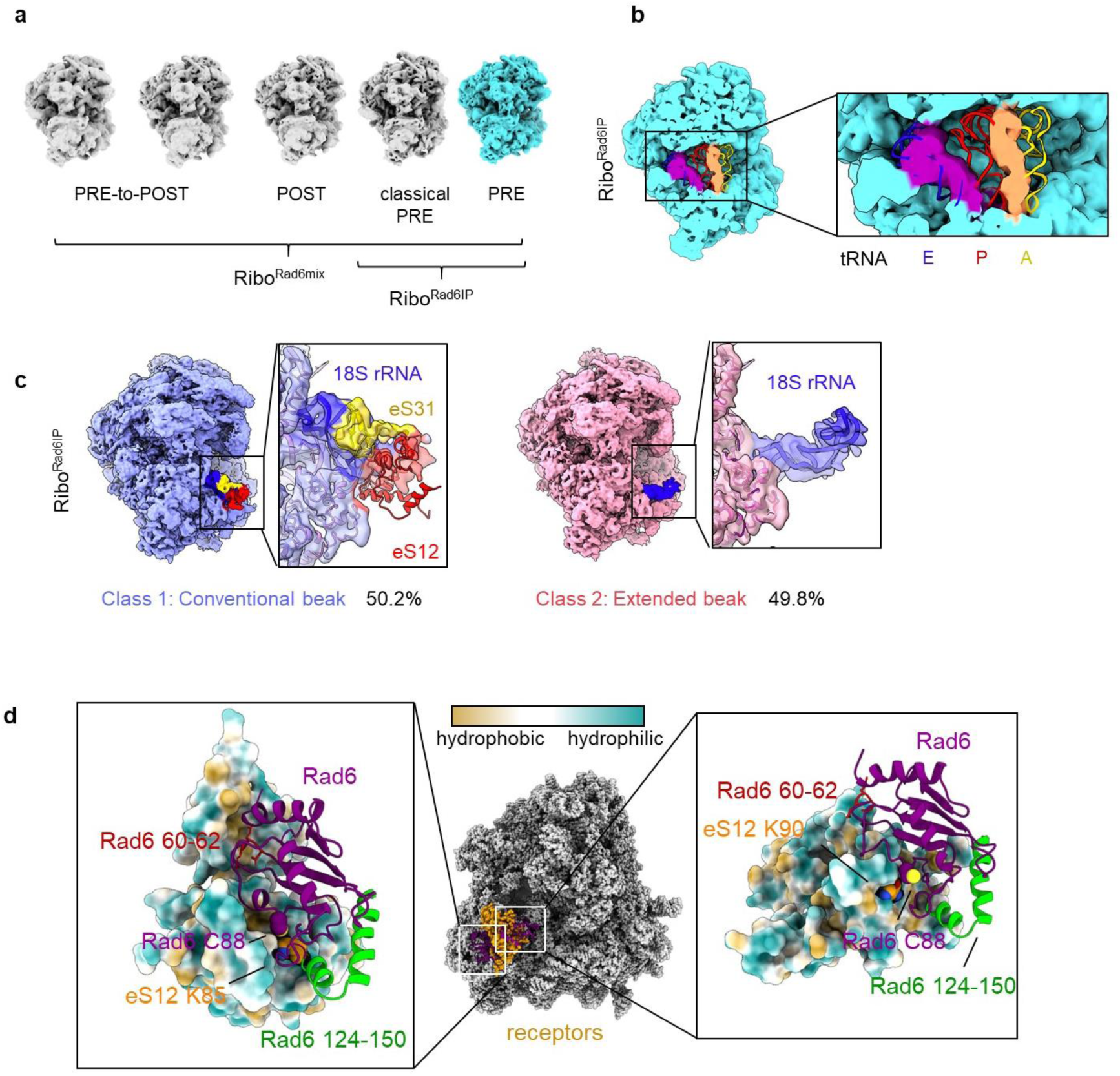
Cryo-EM analysis and molecular modelling of Rad6 binding to ribosomes. **a**, 3D classification of ribosomes incubated with purified Rad6 (Ribo^Rad6mix^) and ribosomes co-immunoprecipitated with FLAG-tagged Rad6 (Ribo^Rad6IP^). Ribosomes were observed in the pre-translocation (PRE), post-translocation (POST), and in the transition state (PRE-to-POST). **b**, Structural detail of Ribo^Rad6IP^ at the pre-translocation stage with tRNAs in hybrid A/P and P/E positions. Rigid fit of the tRNAs was performed using coordinates from the 80S ribosome (PDB ID 6GZ5)^70^. **c,** Focused classification of Ribo^Rad6IP^ shows two classes with distinct conformations for the 40S beak. The extended beak rRNA model was built using COOT^71^. **d,** Molecular docking predicts the binding poses of Rad6 on the ribosome for the two ubiquitination sites on eS12. In the blowouts, the ribosome receptor sites are shown as a surface and colored tan/cyan for hydrophobic/hydrophilic residues. These two binding poses along with those corresponding to eS3, eS20, and eS21 are shown in more detail in Supplementary 7).

### Rad6 is redox regulated by hetero-disulfide formation with Uba1 (E1)

We have previously shown that K63 ubiquitin accumulates in ribosomes because of redox inhibition of the deubiquitinating enzyme, Ubp2^3^. Rad6 is a cysteine-based enzyme that can also be regulated by reactive oxygen species. Following cellular treatment with H_2_O_2_, we observed the formation of a Rad6-containing high molecular weight complex on isolated ribosomes (Fig. 3a and Supplementary Fig. 8a). This complex is reducible by dithiothreitol (DTT), depends on Rad6 catalytic cysteine, and its formation is impaired by high salt concentration and by the cysteine alkylating agent iodoacetamide (IAM), which suggests the formation of a hetero-disulfide (Fig. 3b and Supplementary Fig. 8a,b). We reasoned that in response to stress, Rad6 forms a disulfide complex with proteins with which it closely interacts, such as ribosomal proteins or proteins from the ubiquitination cascade. First, we showed that this disulfide is not formed with a single Rad6-interacting E3 (Supplementary Fig. 8c). Therefore, we performed a denaturing Rad6 co-immunoprecipitation to identify Rad6’s redox partner (Fig. 3c). Using a SILAC-based quantitative mass spectrometry^25^ on the DTT-eluted fraction, we unambiguously determined that Uba1, the yeast E1 activating enzyme, was the major binding partner of Rad6 during H_2_O_2_ stress (Fig. 3d). We validated that E1 was a *bona fide* Rad6 redox partner by showing that Rad6, but not Rad6^C88S^, co-immunoprecipitated with E1 and is released upon DTT reduction (Fig. 3e and Supplementary Fig. 8d). Although other ubiquitin enzymes could interact with E1 and form Cys-based complexes, we showed that in our system, Rad6 is the most abundant E1 partner (Supplementary Fig. 8e). Rad6 mutants with reduced E1 interaction (Rad6^R7A^, Rad6^R11A^, Rad6^R7A/R11A^)^26,27^ do not show growth and ubiquitination defect but are less able to form disulfides in response to stress (Fig. 3f, Supplementary Fig. 8 f-h). Formation of Rad6-containing disulfides depend on the stabilization of a reactive thiolate group in its Cys88. By inspecting the Rad6 catalytic site^21^, we observed three conserved residues able to affect the thiolate stability, N80, Y82, and Q93 (Fig. 3g, Supplementary Fig. 8i). While the Q93E mutation was shown to reduce UBE2A activity by affecting its capacity to deprotonate target lysine residues, a Q93R mutant showed higher ubiquitination activity in *in vitro* reactions^28^. The Q93E mutant partially reduced the accumulation of K63 ubiquitinated targets but the Q93R and N80A mutants severely impaired K63 ubiquitin accumulation and their capacity to form disulfides under stress. Mutations to these residues still allowed these Rad6 variants to be charged by E1 with ubiquitin at different extents (Fig. 3h) but only the N80A mutation severely affected cellular growth (Supplementary Fig. 8j).

**Figure 3.**
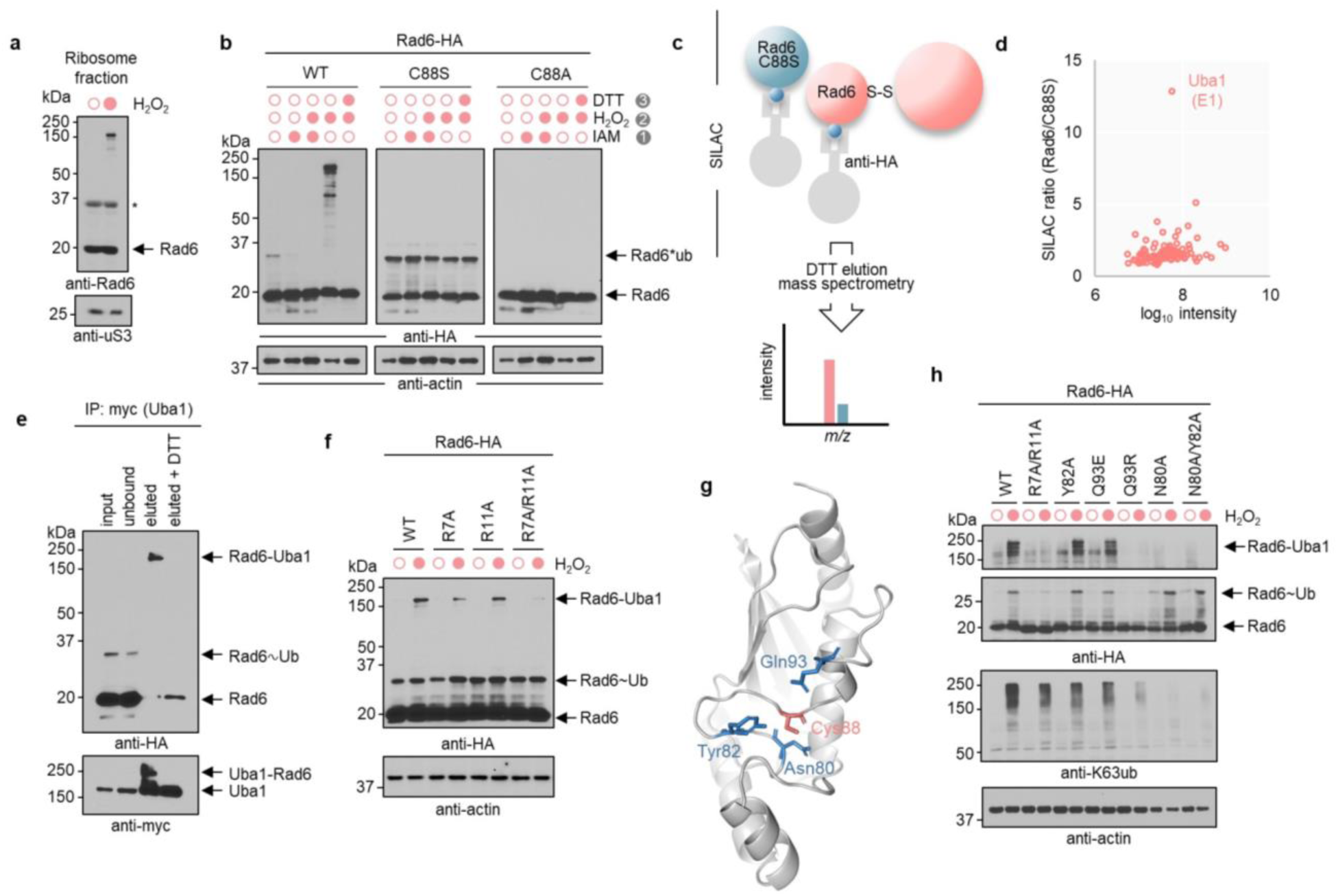
Rad6 is redox regulated by disulfide formation with the E1 enzyme Uba1. **a**, Immunoblot of isolated ribosomes from cells exposed to 0.6 mM H_2_O_2_ for 30 min reveals a high molecular weight complex. * Unspecific band. **b**, Immunoblot of yeast cell lysate expressing HA-tagged wild type Rad6 (WT), C88S, or C88A mutants alkylated with 20 mM IAM, oxidized with 2 mM H_2_O_2_, or reduced with 15 mM DTT. **c**, Schematic of the denaturing co-IP coupled with SILAC LC-MS/MS to identify Rad6 redox partner. **d**, Scatter plot of MS/MS intensity and SILAC ratio of Rad6/C88S under H_2_O_2_ stress. **e**, Immunoblot of Uba1-myc co-IP reveals complex with Rad6-HA, which is reduced by 15 mM DTT. **f**, Rad6 mutants with deficient Uba1 interaction (R7A, R11A, R7A/R11A) show reduced formation of Rad6-Uba1 complex. **g**, Close-up view of Rad6 active site (PDB ID: 1AYZ) highlighting its catalytic cysteine (pink) and vicinal residues (blue) as sticks. **h**, Immunoblots of Rad6 mutants for critical amino acid residues show differential ubiquitination and disulfide formation. Anti-actin was used as loading control.

We observed that oxidative stress leads to the accumulation of K63 ubiquitin while inhibiting Rad6 activity by using its single cysteine residue to form a disulfide complex with the E1 Uba1. Therefore, we postulated that Rad6 would first ubiquitinate ribosomal proteins and subsequently be oxidized and inhibited by disulfide formation. In agreement with this hypothesis, a time course experiment showed that K63 ubiquitin accumulated as early as 2 min of stress induction followed by a subsequent appearance of the Rad6-Uba1 disulfide (Fig. 4a). Importantly, oxidation of Rad6 can occur under oxidation induced by diamide, however, K63 ubiquitin accumulation is specific to H_2_O_2_ (Supplementary Fig. 9a). Next, we observed that disulfide formation depends on the H_2_O_2_ concentration, only a fraction of Rad6 is oxidized under stress, and increased formation of Rad6-Uba1 complex correlates with the depletion of the pool of free ubiquitin (Fig. 4a and Supplementary Fig. 9b). By using mutant strains deleted for DUBs known to decrease the levels of free ubiquitin^29^, we observed increased formation of Rad6-Uba1 disulfides (Supplementary Fig. 9c). Conversely, cells grown to stationary phase, which physiologically contain high amounts of ubiquitin, had increased levels of K63 ubiquitination, no depletion of the pool of free ubiquitin, and no formation of Rad6-Uba1 disulfides *in vitro* or *in vivo,* even at acute 2.5 mM H_2_O_2_ treatment (Fig. 4b).

**Figure 4.**
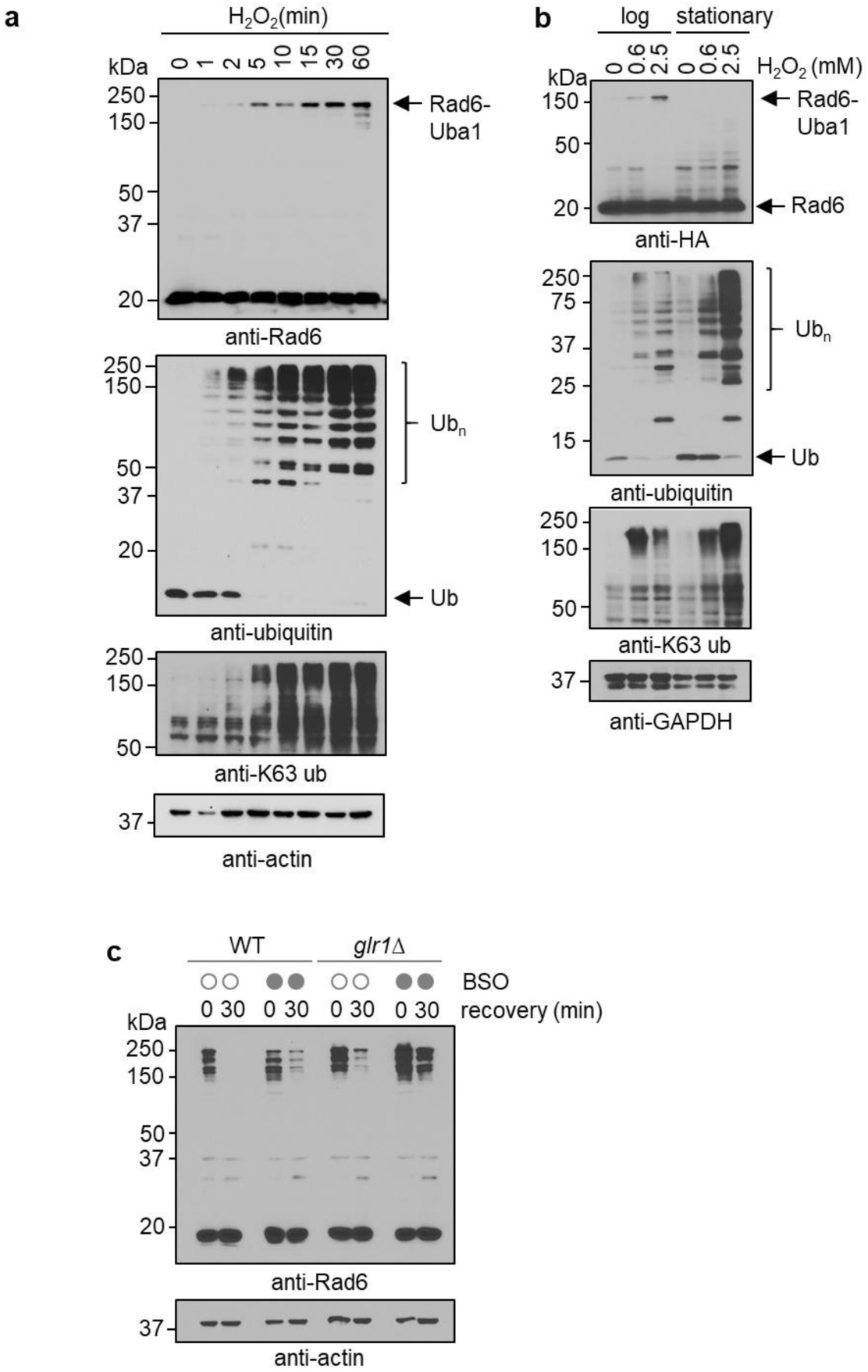
Rad6-Uba1 complex depends on ubiquitin availability and is reduced by the glutathione system. **a**, Rad6-Uba1 and ubiquitin accumulation over time after 0.6 mM H_2_O_2_ treatment. **b**, Immunoblot of cells treated with H_2_O_2_ for 30 min in mid-log or stationary phase. **c,** Reversal of Rad6-Uba1 complex is impacted by the deletion of *GLR1.* Cells were grown into MPD medium^72^ in the presence or absence 2.5 mM of BSO, an inhibitor of glutathione synthesis, for 3 h prior to treatment with 0.6 mM H_2_O_2_ for 30 min and allowed to recover from stress for additional 30 min.

We found that the Rad6-Uba1 complex rapidly disappears during cellular recovery from stress. A pulse-chase experiment showed that Rad6 is not degraded after stress induction, (Supplementary Fig. 9d-e). Inhibition of the proteasome or autophagy did not increase the stability of the Rad6-Uba1 complex (Supplementary Fig. 9f-g), suggesting that thiol reductases are involved in the reduction and recycling of these enzymes. By screening an array of mutants, we showed that deletion of glutathione reductase (*GLR1*) impaired the reduction of Rad6-Uba1 disulfide during stress recovery (Supplementary Fig. 9h-i), while genes from the thioredoxin family had no effect on disulfide reduction (Supplementary Fig. 9j-k). A concomitant treatment with the inhibitor of glutathione synthesis BSO (buthionine sulphoximine) in the *glr1Δ* background strongly prevented the reduction of the disulfide (Fig. 4c). Collectively, our data supports a mechanism by which Rad6-Uba1 is formed following stress induction and is rapidly reversed by glutathione-dependent enzymes once cellular reductive capacity is restored.

We next asked whether this redox regulation of Rad6 and its ability to build K63 ubiquitin chains is evolutionarily conserved in higher eukaryotes. Expression of Rad6 homologs in the *rad6Δ* background have been shown to recover DNA repair functions but not all phenotypes^14,30^. Episomal expression of the human homolog of Rad6 (UBE2A) in the *rad6Δ* background recovers its defective growth phenotype, induces stress resistance, and promotes K63 ubiquitination of ribosomes (Fig. 5a-b and Supplementary Fig. 10a-c). UBE2A also binds to ribosomes (Supplementary Fig. 10c-d) and is able to form disulfides during stress when expressed in yeast and also natively in HeLa cells exposed to H_2_O_2_ (Fig. 5c-d and Supplementary Fig. 10e). Thus, our results suggest that redox ubiquitination of ribosomes can be performed by Rad6 orthologues and that the formation of Rad6-containing disulfide is evolutionarily conserved and regulates UBE2A function in human cells exposed to oxidative stress.

**Figure 5.**
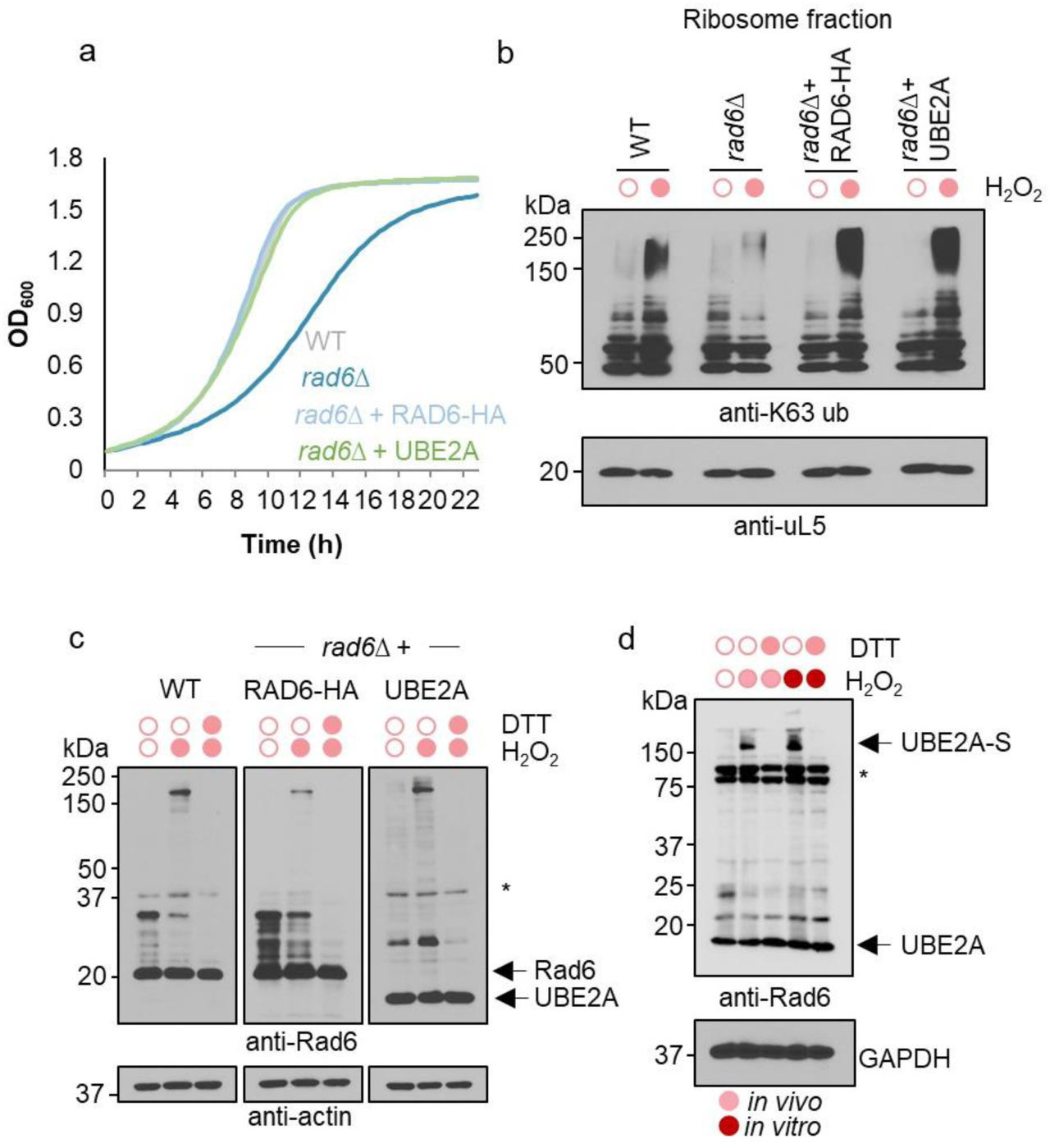
Human UBE2A functionally complements Rad6 in the RTU and is redox regulated. **a**,**b**, Expression of UBE2A reverts *rad6Δ* growth defect (**a**) and ribosomal K63 ubiquitination (**b**). **c**, Wild type and *rad6Δ* yeast cells expressing UBE2A or Rad6-HA form DTT-reducible (15 mM) disulfides under 2 mM H_2_O_2_ incubation *in vitro.* **d**, UBE2A forms disulfides in HeLa cells after *in vivo* (0.5 mM) or *in vitro* (2 mM) incubation with H_2_O_2_, both being reduced by 15 mM DTT. Anti-uL5, actin, and GADPH were used as loading control.

### Rad6 is required for translation reprogramming and activation of integrated stress response

In the RTU, we showed that K63 ubiquitin participates in the regulation of translation elongation in response to oxidative stress^5^. However, the impact of Rad6 in translation and cellular resistance to stress remained underexplored. Similar to the K63 ubiquitin^4^, *RAD6* is necessary for global inhibition of translation that occurs under cellular exposure to H_2_O_2_ (Fig. 6a). Furthermore, WT cells exposed to H_2_O_2_ showed significant inhibition of translation measured by a GFP report system, while the rate of GFP production remained mostly unchanged in *rad6Δ* cells (Fig. 6b). Interestingly, while WT cells reduce their growth upon stress and rapidly resume exponential growth to achieve stationary phase, the growth rate of *rad6Δ* cells is largely unaltered by H_2_O_2_ treatment (Fig. 6c, Supplementary Fig. 11a). FACS analysis showed that H_2_O_2_ treatment led to similar arrest in cell division in wild type and *rad6Δ* cells (Supplementary Fig. 11b), however, increased OD_600_ in *rad6Δ* cells correlates with a prevalence of larger cells (Supplementary Fig. 11c), likely associated with active translation and Rad6-dependent regulation of cyclins^31^. This information prompted us to investigate whether dysregulation of translation could explain why *rad6Δ* cells are more sensitive to H_2_O_2_. First, we showed that *rad6Δ* have higher levels of ROS than WT cells at steady state, which increases significantly after 30 min of stress induction (Fig. 6d). To understand Rad6’s impact on protein production, we employed quantitative mass spectrometry analysis and observed that after normalization by strain differences, only a few proteins were differentially expressed during stress (Fig. 6e,f). This finding suggested that both strains induced the expression of similar antioxidant programs, but the differences are rather quantitative and time-dependent. By inspecting the abundance of antioxidant proteins, we detected unique trends for selective genes involved in detoxifying ROS (Fig. 6f). Thus, we evaluated the levels of antioxidant proteins by immunoblot and determined that Tsa2, Gpx2, and Sod2 are markedly expressed upon stress induction and that this increased protein abundance also lasted longer in *rad6Δ* cells (Fig. 6g). According to our predictions, their protein levels are regulated at the translational level as qPCR analyses revealed similar transcriptional induction in both strains (Fig. 6h). Stalling of translation has been shown to induce the integrated stress response (ISR)^32^, and we hypothesized that reduced translation elongation mediated by K63 ubiquitin would induce the activation of the ISR kinase Gcn2. Indeed, we observed higher levels of phosphorylated initiation factor eIF2α in wild-type cells, which depends on Gcn2 (Fig. 6h-i). Our findings revealed a new pathway of activation of the ISR under cellular exposure to oxidative stress, highlighting a new role for the E2 enzyme Rad6 in the control of translation and resistance to stress.

**Figure 6.**
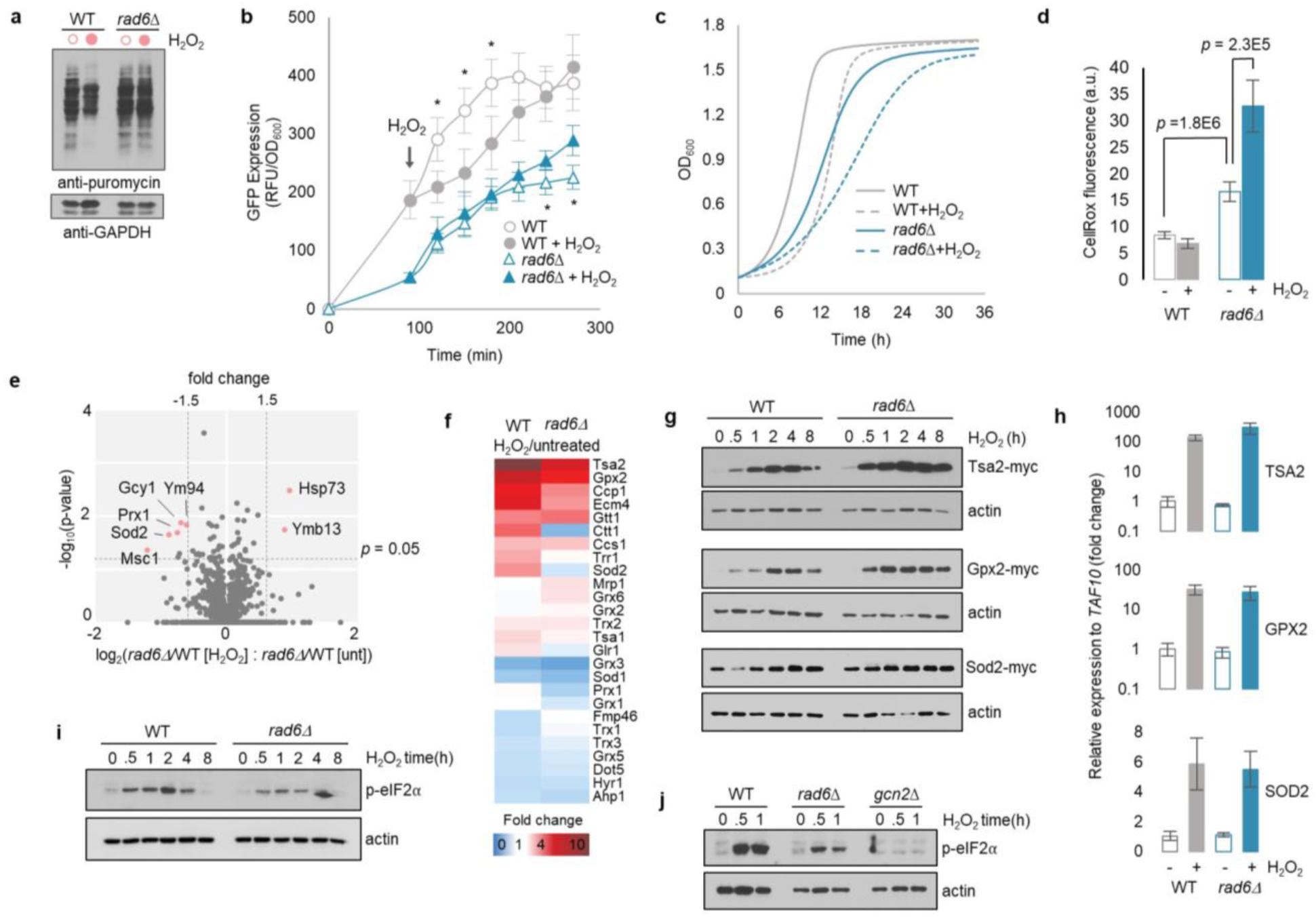
Rad6 supports translation and cellular resistance to stress. **a**, Immunoblot of cellular lysate from wild type (WT) and *rad6Δ* cells in the presence or absence H_2_O_2_ for 15 min, following 30 min of puromycin incorporation (0.9 mM). **b,** Fluorescence determination of GFP in WT and *rad6Δ* cells. GFP expression was under a Met25 promoter and was induced at time zero by transferring cells to Met-depleted medium. * p<0.05. **c**, Growth curve of WT and *rad6Δ* strains in the presence (dashed) or absence (solid) H_2_O_2_. **d,** ROS determination by 2.5 µM CellRox Deep Red fluorometric assay (Thermo Scientific) in lysates from WT and *rad6Δ* cells. Fluorescence was captured at 665 nm with excitation at 644 nm. **e**, Volcano plot of SILAC proteomics ratio from H_2_O-treated and untreated (unt) WT and *rad6Δ* cells. **f**, Heatmap of MS/MS intensity fold change for antioxidant proteins in the WT and *rad6Δ* cells. **g,** Immunoblot anti-myc for the expression of the antioxidant proteins Tsa2, Gpx2, and Sod2 in WT and *rad6Δ* cells. **h,** Bar graph of qPCR analysis of *TSA2*, *GPX2*, and *SOD2* transcripts levels relative to TAF10 from untreated and H_2_O_2_-treated (30 min) WT and *rad6Δ* cells. **i** Immunoblot anti-phosphorylated eIF2α (p-eIF2α) in the WT and *rad6Δ* cells. **j,** Immunoblot anti-p-eIF2α in the WT, *rad6Δ*, and *gcn2Δ* cells. Anti-GAPDH and anti-actin were used as loading control.

## Discussion

Our work characterized a new function for the ubiquitin conjugating enzyme Rad6 in the regulation of cellular response to oxidative stress in the pathway of Redox control of Translation by Ubiquitin (RTU). In this pathway, we showed that Rad6 binds (Fig. 1d,e) and modifies ribosomes with K63 polyubiquitin chains under stress (Fig. 1b,c). Using single-particle cryo-EM, we showed that Rad6 interacts with ribosomes at the pre-translocation stage of elongation, highlighting new structural features in the 40S beak (Fig. 2) related to either binding or stabilization of Rad6 after recruitment, as predicted by our docking calculations (Fig. 2d). Although Ubc13 is the main E2 enzyme known to be involved in K63 ubiquitination^13^, yeast cells lacking *UBC13* and ribosomes isolated from *ubc13Δ* strain were still modified by K63 ubiquitin in the presence of Rad6 and H_2_O_2_ (Supplementary Fig. 1a, 2e). In the RQC (Ribosome-associated Quality Control) and NRD (non-functional rRNA decay) pathways^33,34^, Ubc4 has been suggested as the E2 associated with the E3 Hel2 to promote ribosome ubiquitination^20,35^. However, deletion of both paralogs *UBC4* and *UBC5* did not prevent K63 ubiquitin accumulation under stress^3^, nor did the deletion of *HEL2* (Supplementary Fig. 1b). Thus, our results suggest that Rad6 modifies ribosomes in the RTU independent of other pathways of translation control, suggesting a potential functional code for ribosome ubiquitination. It is likely that these pathways occur concomitantly in a complex and dynamic system such as the cellular response to stress, targeting different subpopulation of ribosomes, with Rad6 carrying out most of the K63 ubiquitination load during oxidative stress.

Our working model proposes that inhibition of the DUB Ubp2 by H_2_O_2_ is the first event of the RTU that acts as a stress sensor and leads to the accumulation of K63 ubiquitinated ribosomes conjugate by Rad6. This model assumes that ubiquitination and deubiquitination of ribosomes are regular components of the translation cycle, and K63 ubiquitin accumulates when deubiquitination is impaired. This model is supported by the constitutive presence of Rad6 on ribosomes and polysomes (Fig. 1e) and by the fact that increased translation initiation leads to higher levels of K63 ubiquitin even in the absence of stress^3^. Moreover, the lack of K63 ubiquitin impacts polysome stability and renders cells more susceptible to translation inhibitors in the absence of stress^3,36^. Molecular details of the regulatory role of K63 ubiquitin in the translation cycle are not fully understood and our future studies will focus on defining how basal levels of K63 ubiquitin impact translation and why this modification is required to increase the stability of polysomes.

Several enzymes have been shown to be modified by ROS, acting as a redox switch to rapidly regulate their activity^37,38^. Although many enzymes regulated by ROS contain catalytic cysteine residues in their active sites, their susceptibility to oxidation requires a structural organization that renders a reactive thiolate group^39^ and a conducive cellular environment that fosters the formation of oxidized species. For example, it has been suggested that only a fraction of DUBs from the cysteine-containing USP family, can be redox regulated and re-activated by DTT^9^. Previous work had proposed that the yeast Cdc34 would be the only E2 oxidized to form disulfides during oxidative stress induced by diamide or H_2_O_2_ in rich medium^40^. Here we showed that Rad6 is redox-regulated (Fig. 3b) and is the main protein forming disulfides with the E1 Uba1 during peroxide stress in our system (Fig. 3e,f). We also showed that the human homolog of Rad6 (UBE2A) complements Rad6 function and is redox regulated in yeast and in HeLa cells (Fig. 5b-d). Considering Rad6 structural features, we demonstrated that residues that foster Rad6 interaction with the E1 (R7 and R11) and a conserved asparagine residue (N80) are important for Rad6 activity and its capacity to form disulfides (Fig. 3f,h). The human SUMO E2 (Ubc9) also forms disulfides with E1 during oxidative stress and incapacity to maintain these disulfide leads to cellular sensitive to stress^41,42^. Distinct from Rad6-mediated accumulation of K63 ubiquitin, Ubc9 oxidation abolishes protein SUMOylation revealing important mechanistic differences despite sharing a similar chemical regulatory process, which highlights the need to investigate each redox pathway individually.

Besides the structural susceptibility, disulfide formation also depends on the pool of free ubiquitin, Rad6 subcellular location, and the availability of the E1 enzyme. Our data suggest that Rad6 interaction with uncharged E1 would favor a conformation that exposes their cysteine residues and fosters the formation of a disulfide bond in the presence of H_2_O_2_. Furthermore, we showed that the Rad6-Uba1 complex is not degraded during recovery from stress and requires the glutathione system to be reversed in the cellular context (Fig. 4c and Supplementary Fig. 9d-i). We speculate that formation of this Rad6-Uba1 disulfide protects these enzymes from hyperoxidation and the reversibility of this process would allow cells to quickly recycle and make them readily available to ubiquitinate substrates. We further propose that Rad6 redox regulation by disulfide formation could serve as a mechanism to curb the levels of K63 ubiquitin and fine tune translation. This mechanism would also prevent a complete depletion of the pool of free ubiquitin, given the abundance of ribosomes.

In the RTU, K63 ubiquitin is required to inhibit translation during cellular exposure to stress^3,4^. K63 ubiquitin modifies fully assembled 80S and polysomes and these complexes were mostly present in the pre-translocation stage of translation elongation, with a marked destabilization of the 60S P-stalk when compared to ribosomes from the K63R ubiquitin mutant strain^5^. Here we showed that *RAD6* is required for global inhibition of translation during cellular exposure to H_2_O_2_ (Fig. 6a,b). We reasoned that dysregulation of translation by the absence of K63 ubiquitinated ribosomes led to higher levels of ROS and increased sensitivity to stress (Fig. 6c,d). In agreement with this hypothesis, *rad6Δ* showed increased levels of protein production, including antioxidant enzymes (Fig. 6a,b,g). Deletion of *RAD6* also inhibited the phosphorylation of the translation initiation factor eIF2α and impacted the synthesis of the stress response transcription factor Gcn4 (Fig. 6i,j). The integrated stress response starts with activation of the Gcn2 kinase causing phosphorylation of the translation initiation factor eIF2α^43^. Phosphorylation of eIF2α (p-eIF2α) suppresses the formation of ternary complex (eIF2–GTP–Met-tRNA_i_^Met^), which leads to reduced translation initiation while fostering the synthesis of stress-related proteins^43^. Activation of Gcn2 was classically defined by its binding to deacylated tRNAs^44–46^. Since Rad6 is important for DNA repair^18,47^, we initially discussed the possibility that extensive nucleic acid damage in response to oxidative stress would trigger the ISR as a result of induced deacylation of tRNAs and activation of quality control pathways^48^. However, different from the RQC that antagonizes the ISR during DNA damage^49,50^, we showed that Rad6 activity is required to activate ISR and increase the levels of p-eIF2α induced by Gcn2 (Fig. 6j), likely through stalling of translation at elongation. An alternative model of activation of Gcn2 has been proposed in recent studies through its binding to ribosomes^51–56^. Furthermore, it has been shown that translation stalling and Gcn2 binding to proteins in the 60S P-stalk not engaged with elongation factors triggers the ISR^32,53,54^. Therefore, we propose that destabilization of the 60S P-stalk by K63 ubiquitin would provide a cross-talk mechanism between translation elongation and initiation under stress^5^. As a potential redox switch in the RTU, it will be also essential to understand the mechanisms of recruitment and activation of Gcn2 during oxidative stress as well as the fate of deubiquitinated ribosomes.

Finally, Rad6 is a highly conserved and multi-functional protein, whose mutations in higher eukaryotes have been associated with cognitive and learning deficits^16^, and in humans have led to the intellectual disability type Nascimento^57^ and the progression of chronic myeloid leukemia^58^. UBE2A also binds the E3 ligase Parkin, which ubiquitinates proteins involved in the clearance of defective mitochondria^59^. We have shown that disruption of Rad6 interaction with E1 (Rad6^R7A/R11A^) impairs its capacity to form disulfides under stress (Fig. 3g) and mutations to UBE2A Arg7 have been identified in individuals with the Nascimento syndrome^15^, highlighting the impact of a UBE2A regulatory mutations on patient’s cognitive development. Previous studies have shown that there is a spectrum of clinical phenotypes for the Nascimento disability depending on the UBE2A mutation carried by these individuals^15^. The different domains that control UBE2A activity, protein interaction, translation control, and activation of the integrated stress response could provide important ways to explain the range of observed clinical phenotypes. Defining the role of UBE2A domains, their function in translation reprogramming, and their effect in stress response will be key to understanding the development and resistance against stress-related diseases.

## Acknowledgements

We thank Matt Foster and the Duke Proteomics and Metabolomics Shared Resource for the support with mass spectrometry data acquisition. We are indebted to Nicholas Brown for a kind donation of ubiquitin constructs. We also thank Paul Magwene, Masayuki Onishi, and Debra Murray for making equipment available to this project. This work was supported by US National Institutes of Health R00 Award ES025835 and R35 Award GM137954 (G.M.S.). This work was supported in part by Intramural Research Program of the NIH; National Institute of Environmental Health Sciences Grant ZIC ES103326 (M.J.B.). Cryo-EM work was performed at the Duke University Shared Materials Instrumentation Facility (SMIF), a member of the North Carolina Research Triangle Nanotechnology Network (RTNN), which is supported by the National Science Foundation (Grant ECCS-1542015) as part of the National Nanotechnology Coordinated Infrastructure (NNCI). Because of COVID19 restrictions, cryo-EM grids were also prepared and screened at the UNC Chapel Hill CryoEM Core Facility. We thank Joshua Strauss for his technical assistance. We also acknowledge UNC Lineberger Comprehensive Cancer Center through the University Cancer Research Fund and the Cancer Center Support Grant P30CA016086.

## Author contributions

G.M.S. conceived and funded the project. V.S. and G.M.S. designed experiments and wrote the manuscript. All authors conducted experiments, data analysis, edited, and commented on the manuscript.

## Materials and Correspondence

Requests and correspondence should be addressed to corresponding author Gustavo Silva

## Data Availability

Two Ribo^Rad6IP^ EM maps were deposited in the Electron Microscopy Data Bank (EMDB), https://www.ebi.ac.uk/pdbe/emdb. The mass spectrometry proteomics data have been deposited to the ProteomeXchange Consortium via the PRIDE partner repository.

## Methods

### Cell strains and culture

All yeast *Saccharomyces cerevisiae* strains used in this study are described in Supplementary Table 3 and all plasmids are listed in Supplementary Table 4. Standard recombination methods were used to delete and tag genes; deletions were confirmed by PCR and tagged genes were confirmed by PCR and immunoblotting. Unless specified, cells were cultivated into synthetic dextrose minimal medium (SD: 0.67% yeast nitrogen base, 2% dextrose and required amino acids) at 30 °C at 200 rpm agitation. Cells were disrupted by glass-bead agitation at 4 °C in standard buffer: 50 mM Tris-HCl pH 7.5, 150 mM NaCl, 20 mM iodoacetamide (IAM), 1X EMD Millipore protease inhibitor cocktail set I. The extract was cleared by centrifugation, and protein concentration was determined by Bradford assay (Bio-Rad).

### Growth assays

Yeast cultures were grown into SD medium to mid-log phase and back-diluted to OD_600_ 0.1. Equal volumes of cell suspension and fresh SD medium (75 µl) were added to a 96-well plate in triplicate. Plates were incubated at 30 °C with shaking, and absorbance was measured at 595 nm every 15 min over a period of 24 - 48 h in a Tecan Sunrise microplate reader. For sensitivity assays, indicated strains were grown into SD medium to mid-log phase and back-diluted to OD_600_ of 0.1. Serial five-fold dilutions were spotted onto YPD plates and incubated for 2 days at 30 °C.

### Mammalian cell culture

HeLa cells (ATCC CCL2) were cultured in EMEM (ATCC 30-2003) containing 10 % fetal bovine serum (Invitrogen 16140071) and 1 % Penicillin/Streptomycin (Invitrogen 15140122). Cells were treated with 500 μM H_2_O_2_ at 70 % confluence one day after being seeded. Protein extraction was performed by cell sonication in standard buffer containing 0.5 % NP40. The extract was cleared by centrifugation, and protein concentration was determined by Bradford assay (Bio-Rad) prior to western blotting.

### Protein purification

Rad6 and Bre1 were expressed in pGEX vector containing GST-tag and TEV protease sequences. Expression of yeast Rad6 and Bre1 was induced in *E. coli* BL21-CodonPlus (DE3)-RIL with 0.6 mM isopropyl thio-β-galactopyranoside (IPTG) for 4 h at 37 °C or overnight at 16 °C, respectively. Cells were lysed by sonication in NETN buffer (50 mM Tris-HCl pH 7.5, 150 mM NaCl, 0.5% IGEPAL) with 0.15 mM DTT, 1 mM PMSF, and lysozyme (10mg/ml). Lysate was cleared by centrifugation at 12,000 x g for 20 min at 4° C. Glutathione-containing beads (Goldbio G-250-5) were pre-washed with NETN buffer (3x) and incubated for 2 h at 4 °C under rotation with the cleared lysate. Beads were washed twice with NETN buffer followed by two washes with 1X PBS buffer. Proteins were eluted by cleavage of the GST tag by 15 units/ml TEV protease (Sigma T4455-1KU) in PBS buffer overnight at 4 °C. The eluted proteins were desalted using a PD-10 column in 50 mM Tris-HCl pH 7.5, 100 mM NaCl buffer. Purity was evaluated by SDS-PAGE. Rad6 activity was determined by its charging capacity, which was visualized by immunoblot. Purified Rad6 (500 nM) was incubated with E1 (21 nM), ubiquitin (1.25 µM), and ATP (10 mM) in 20 mM HEPES buffer pH 7.2 containing 10 mM MgCl_2_ and 1 mM DTT for 30 min at 30 °C.

### Ubiquitination assay

Ubiquitination reactions were performed in the presence of 100 nM E1 UBA1 (Enzo Life Sciences BML-PW8395-0025), 250 nM Rad6, 150 nM of Bre1, 2-10 µM of ubiquitin (LifeSensors si201), 10 µg of isolated ribosomes, and energy regenerating system (ERS – 1 mM ATP, 10 mM creatine phosphate, 20 µg/ml creatine kinase). All components were pre-incubated in reaction buffer (50 mM Tris-HCl pH 7.5, 150 mM NaCl, 2 mM MgCl_2_, 1 mM dithiothreitol) for 10 min at room temperature, before the addition of Bre1 and ribosomes. The reaction was incubated for up to 3 hours at 30 °C at 300 x rpm agitation and stopped by the addition of Laemmli buffer. When indicated, samples were incubated with 20 mM DTT for 10 min prior to western blotting.

### Western blotting

Proteins were separated by standard 10 % - 15 % SDS-PAGE loaded in Laemmli buffer. Samples were transferred to PVDF membrane (ThermoFisher), and immunoblotting was performed using the following antibodies: anti-K63 ubiquitin (1:4,000; EMD Millipore, cat. no. 05-1308, clone apu3), anti-actin (1:6,000; Cell Signaling, cat. no. 4967), anti-GAPDH (1:4,000; Abcam, cat. no. ab9485), anti-Myc (1:5,000; ThermoFisher, cat. no. R950-25 and PA1-981), anti-Rad6 (1:8,000; Abcam, cat. no. ab31917), anti-Rps3/uS3 (1:6,000; Cell Signaling, cat. no. 9538S), anti-Rpl11/uL5 (1:6,000; Cell Signaling, cat. no. 18163S), anti-HA (1:10,000; ThermoFisher, cat. no. 71-5500), anti-Rps20/uS10 (1:9,000; ThermoFisher, cat. no. PA5-75383), anti-ubiquitin (1:10,000; Cell Signaling Technology cat. No. 3936S), anti-FLAG (1:3,000; MilliporeSigma, cat. no. C986X12), anti-puromycin (1:4,000; MilliporeSigma, cat. no. MABE343). Anti-mouse (1:8,000-12,000; Abcam, cat. no. ab97245) and anti-rabbit (1:6,000-12,000; GE Healthcare – Life Sciences, ca. no. NA934) secondary antibodies conjugated with HRP and ECL Prime detection reagents were acquired from GE Healthcare Life Sciences. All antibodies have been validated by the manufacturer or are expected to react with the species used in this study based on sequence similarity.

### Sucrose sedimentation profile

Yeast cells were incubated in the presence of 150 μg/ml cycloheximide for 5 min at 30 °C and lysed immediately in extraction buffer (20 mM Tris-acetate, pH 7.0, 50 mM NaCl, 15 mM MgCl_2_, 20 mM IAM, 200 μg/ml heparin, 150 μg/ml CHX, 1x complete mini EDTA-free protease inhibitor cocktail (Roche). A total of 400 μg of RNA was sedimented by ultracentrifugation for 150 min at 36,000 rpm (Beckman SW40 rotor) at 4 °C in a 7–47 % sucrose gradient (buffered in 50 mM Tris-acetate, pH 7.0, 15 mM MgCl_2_, 150 μg/ml CHX). During sucrose gradient analysis, absorbance was monitored at 254 nm while fractions were collected using a Brandel density gradient fractionation system equipped with the PeakChart software. Proteins from fractions were precipitated with TCA-acetone prior to western blotting.

### Ribosome isolation

Yeast cells were lysed as for the sedimentation profile. A total of 400 μg of RNA (determined by A_260_) was sedimented by ultracentrifugation for 120 min at 70,000 rpm (Beckman Optima Max-TL, TLA-110 rotor) at 4 °C in a 50% sucrose cushion buffered in 50 mM Tris-acetate, pH 7.0, 150 mM NaCl, and 15 mM MgCl_2_. Ribosome pellet was resuspended in 50 mM Tris-acetate, pH 7.0, 150 mM NaCl and ribosome-free supernatant was precipitated with TCA-acetone, followed by resuspension to the same final volume prior to immunoblotting.

### Cryo-electron microscopy analysis

#### Sample preparation

1*) Ribosome isolation for in vitro incubation with Rad6 (Ribo^Rad6mix^)*: Yeast cell lysate from *rad6Δ* strain was prepared as above and ribosome pellets were resuspended in buffer containing 50 mM Tris-Acetate pH 7.0, 150 mM NaCl, 30 mM MgCl_2_. 20 µg of ribosomes were incubated with 1.2 µM of purified Rad6. *2) Ribosome co-immunoprecipitation with Rad6-FLAG (Ribo^Rad6IP^)*: Yeast cell lysate from Rad6-FLAG strain (GMS488) was cleared by centrifugation and loaded into a column containing M2 anti-FLAG resin (Sigma-Aldrich). The sample was eluted with 100 μg/ml of 3X FLAG peptide (Sigma-Aldrich) in buffer containing 50 mM Tris-HCl pH 7.5, 100 mM NaCl, 5 mM MgCl_2_, and 10% glycerol. Enrichment for large complexes and sample concentration was performed using Amicon centrifugal filter units with 100 kDa cutoff. Rad6-FLAG bound ribosomes were confirmed by immunoblot prior to cryo-EM analysis. *3) Cross-linking of ribosomes co-immunoprecipitated with Rad6-FLAG (Ribo^Rad6IP-XL^)*: Ribo^Rad6IP^ prepared as above were incubated with 500 µM BS^3^ (ThermoFisher) for 2h at 4 °C.

#### Grid preparation

For Ribo^Rad6IP^ and Ribo^Rad6IP-XL^, 3 µl of the samples at 1 µg/µl was applied to UltrAUfoil R1.2/1.3 grids (Quantifoil Micro Tools GmbH). Grids were glow-discharged for 30 sec at 15 mA prior to sample application (PELCO easiGlow™, Ted Pella, Inc.). Grids were blotted between 2.5 and 3 sec and plunged frozen in liquid ethane using a Leica vitrification plunger (EM GP2, Leica Microsystems). For Ribo^Rad6mix^, cryo-grids were prepared by rapid immersion into liquid ethane with a Vitrobot Mark IV (Thermo Fisher Scientific) set to 22 °C and 95% humidity. Quantifoil 300 mesh R1.2/1.3 (Quantifoil) grids were used and glow-discharged as above.

#### Data collection

Ribo^Rad6mix^ and Ribo^Rad6IP^ micrographs were recorded under low-dose conditions on a FEI Titan Krios electron microscope (ThermoFisher Scientific) equipped with a K3 direct electron detector using the Latitude software (Gatan, Pleasanton, CA). A single hole was targeted per stage movement, and one shot was used per hole. Sixty frames at a nominal pixel size of 1.08 Å were collected per movie. The cumulative electron dose was 60 e^−^/Å^2^, and 3,062 movies and 7,872 movies were collected for Ribo^Rad6IP^ and Ribo^Rad6mix^, respectively. Ribo^Rad6IP-XL^ data were collected using SerialEM ^60^ on a Talos Arctica microscope (Thermo Fisher) equipped with a K2 detector. Data was collected at a magnification of 45,000x (0.932 Å /pix). Movies were collected using a dose rate of ∼0.8 e^−^/Å^2^/frame and a total of 60 frames were acquired over an 8.4 sec exposure. A total of 3,372 movies were collected using 9 beam-image shift exposures per stage movement (3 x 3 hole pattern).

#### Single-Particle Data Processing

For Ribo^Rad6mix^, movie alignment was performed in RELION^61^, and the contrast transfer function was determined on the motion-corrected sum of frames using CTFFIND4.1^62^. Micrographs were imported into cryoSPARC^63^ for particle picking and 2D classification. Around 1 M particles were selected and re-extracted with a binning factor of 4 (pixel size 4.32 Å) in RELION for further processing. Another round of 2D classification was done in RELION resulting in 651,713 particles which were subjected to 3D classification without masking. After removing the noise classes, the remaining 458,977 particles were re-extracted using a binning factor of 2 (pixel size 2.16 Å) and subjected to 3D refinement resulting in a map of 4.4 Å resolution. Polishing was performed to correct for the local motion of all refined particles. At this point, two different strategies were employed. First, we carried out focused classification to analyze the conformational changes of the 40S beak. All polished particles were refined with a shape mask only including the 40S domain. An elliptical shape mask located at the 40S beak region was then used for focused classification into six classes (Supplementary Fig.6a). All classes were selected and used for 3D reconstruction without alignment to visualize the overall ribosome structure. For global classification, all polished particles were imported into *cis*TEM^64^ and local 3D classification was performed using the refinement parameters obtained from RELION. Four out of the six classes showed well-defined tRNA features.

For the Ribo^Rad6IP^ dataset, all the data processing was done in RELION. A total of 245,077 particles were picked and extracted with a pixel size 1.8 Å. Next, 224,217 particles were selected after 2D classification and subjected to 3D classification without masking. Three good classes consisting of 68,538 particles were selected and refined to 4.5 Å. Further 3D classification without alignment resulted in two different conformations of ribosomes (rotated PRE translocation state and classical PRE translocation state) and one noise class. For the 40S focused classification, all good 3D classes were pooled and re-extracted using a pixel size of 1.08 Å. Then, 3D refinement with a 40S shape mask was performed to align exclusively the 40S domain. Another round of 3D classification was conducted with an elliptical shape mask focusing on the 40S beak yielding two different conformations of the focus region (conventional and extended 18S rRNA). All classes were used to perform a consensus 3D reconstruction showing the overall ribosome features. Two classes showing the same conformation were combined (31,644 particles) to obtain the final map. No further refinement was conducted to avoid overfitting of the 40S beak region due to its small size.

For Ribo^Rad6IP-XL^ dataset, 2,638 aligned micrographs were imported into cryoSPARC. CTF estimation was conducted with CTFFIND4.1. A total of 1,514 micrographs were selected based on the CTF maximum resolution using a cutoff of 4.5 Å, from which 105,705 particles were selected using a blob picking job. After 2D classification, a total of 31,670 particles were selected and subjected to Ab-initio reconstruction. Final homogenous refinement resulted in 4.4 Å resolution map.

### Molecular Modeling

The coordinates for Rad6 were taken from the crystal structure of Rad6 with ubiquitin (PDB: 4R62)^21^. Ubiquitin was removed from the structure and the mutant C88K was reverted to wild type C88 by replacing the sidechain from the Dunbrack Rotamer Library^65^. The coordinates for the receptor ribosome were taken from a cryo-EM reconstruction of the 80S ribosome in complex with mRNA, tRNA, and eEF2 (PDB: 6GQ1)^66^. Missing sidechains were built in with the Dunbrack Rotamer Library. Receptor sites (such as eS12 and eS21) along with neighboring proteins, were isolated from the 80S ribosome prior to docking to limit search space. Rad6 was then docked to each receptor site using ClusPro 2.0 ^67^. To further refine our search, we added attraction benefits to Rad6 residues 60-139, encompassing residues characterized in this study known to be important for binding. On each receptor site we added attraction benefits to the putative ubiquitinated lysine residues^4^; no restraints were added that specifically promote Rad6 C88 to interact with these lysine residues. To discriminate between the top ten scoring predictions from ClusPro 2.0 ^68^, we further considered whether Rad6 C88 is proximal to the putative ubiquitinated lysine to promote catalysis.

### Mass Spectrometry analysis

#### Sample preparation

Yeast cells were cultured to mid-log phase into synthetic SILAC media supplemented with light or heavy lysine isotopes (L-Lys8 ^13^C, ^15^N; Cambridge Isotopes). Cells were grown for at least 10 generations in SILAC medium before being treated with 0.6 mM H_2_O_2_ for 30 min at 30 °C. *Rad6 immunoprecipitation*: Rad6-HA (light, GMS483) and Rad6^C88S^-HA (heavy, GMS493) cells were disrupted under denaturing conditions in standard buffer containing 8M urea and 5 mM EDTA. Urea was diluted to 1M prior to anti-HA immunoprecipitation (2 h at 4 °C) and beads were washed for 5 times in 50 mM ammonium bicarbonate pH 8.0. Samples were eluted by 30 min incubation with 15 mM DTT, digested overnight with Trypsin/Lys-C (1:20 w/w, Promega), desalted using C18 Hypersep Spin Column (Thermo), and resuspended in 2% acetonitrile (MeCN), 0.1% formic acid. *Protein expression changes* (*rad6Δ*/WT): WT (GMS280) was grown under SILAC light condition and *rad6Δ* (GMS494) under heavy. Cell lysis was performed as above following trypsin digestion and C18 clean up. Biological triplicate were run combining equal protein amounts per strain from untreated and H_2_O_2_-treated conditions.

#### Data collection & analysis

Up to 1 µg of digests were analyzed using a nanoACQUITY UPLC system (Waters) coupled to a Fusion Lumos high-resolution accurate mass tandem mass spectrometer (ThermoFisher Scientific) via a nano-electrospray flex ion source. Briefly, peptides were trapped on a Symmetry C18 180 μm × 20 mm column (5 μl/min at 0.1 % MeCN) followed by an analytical separation using a 1.7-μm ACQUITY UPLC HSS T3 C18 75 μm × 250 mm column (Waters) with a 90-minute gradient of 5–30 % MeCN with 0.1% formic acid at a flow rate of 400 nl/min and column temperature of 55°C. Data collection on the Fusion Lumos MS was performed in data-dependent acquisition (DDA) mode with a 120,000-resolution (at *m/z* 200) full MS scan from 375-1500 *m/z* with a target automatic gain control (AGC) value of 4e5 ions and maximum IT of 50 ms. Peptides were selected for data-dependent MS/MS using charge state filtering, monoisotopic precursor selection, and a dynamic exclusion of 20 sec. MS/MS was performed using higher energy C-Trap dissociation with a collision energy of 30 ± 5 % with detection in the ion trap using rapid scanning, an AGC target of 1e4 and maximum IT of 100 ms. A time-dependent (2 sec) method was used. The RAW data files were processed using MaxQuant to identify and quantify protein and peptide abundance. The spectra were matched against the yeast *S. cerevisiae* Uniprot database. Protein identification was performed using 10 ppm tolerance with a posterior global FDR of 1% based on the reverse sequence of the yeast FASTA file. Up to two missed trypsin cleavages were allowed, and oxidation of methionine and N-terminal acetylation were searched as variable post-translational modification while cysteine carbamidomethylation was searched as fixed.

#### GFP expression determination

Indicated strains were grown in SD-Ura to mid log phase. At OD_600_ 0.3-0.4, cells were pelleted and resuspended in SD-Ura-Met medium to induce expression of GFP under control of the Met25 promoter. After one hour of growth in Met-deficient medium, the cultures were incubated in the presence or absence of 0.6mM H_2_O_2_. At the indicated time points, cells were collected by centrifugation, washed, and resuspended in PBS for determination of OD_600_ and fluorescence. Data was normalized to the relative fluorescence unit (RFU) of GFP per OD_600._

### Quantitative RT-PCR

Total RNA from yeast cells grown into SD-Trp-Leu to mid-log phase was isolated with YeaStar RNA Kit (Zymo Research R1002). RNA was treated with 0.02 units/µl DNase I (NEB M0303) for 10 min at 37 °C, purified with RNA clean up kit (NEB T2050), and cDNA was generated with SuperScript III Reverse Transcriptase (Thermo Scientific 18080044) from 2-3 µg of RNA using oligo(dT) (Invitrogen 18418012) as primer. Quantitative RT-PCR was performed with 20 ng cDNA in a LightCycler instrument (Roche) using FastStart Essential DNA Green Master (Roche 06402712001). Primers for quantitative RT-PCR are listed in Supplementary Table 5. The relative transcript abundance was normalized to the expression levels of the TAF10 gene. The relative fold change was obtained by ΔΔCt method^69^. Abundance was determined using three biological replicates.

### Flow cytometry

WT and *rad6Δ* strains were grown to mid-log phase and 500 µl of untreated and H_2_O_2_-treated cells were fixed in 70% ethanol overnight at 4 °C. Cells were centrifuged and incubated with 40 μg/ml RNAse A (ThermoFisher EN0531) in 50 mM Na-citrate buffer for 16 h at 50 °C, followed by incubation with 100 μg/ml proteinase K (NEB P8107) for 16 h at 50 °C. Cells were sonicated for 5 sec and SYTOX Green (Invitrogen S7020) at 2 μM was added to the cells before analysis in a BD FACSCanto cell sorter.

---

**Supplementary Figure 1.**
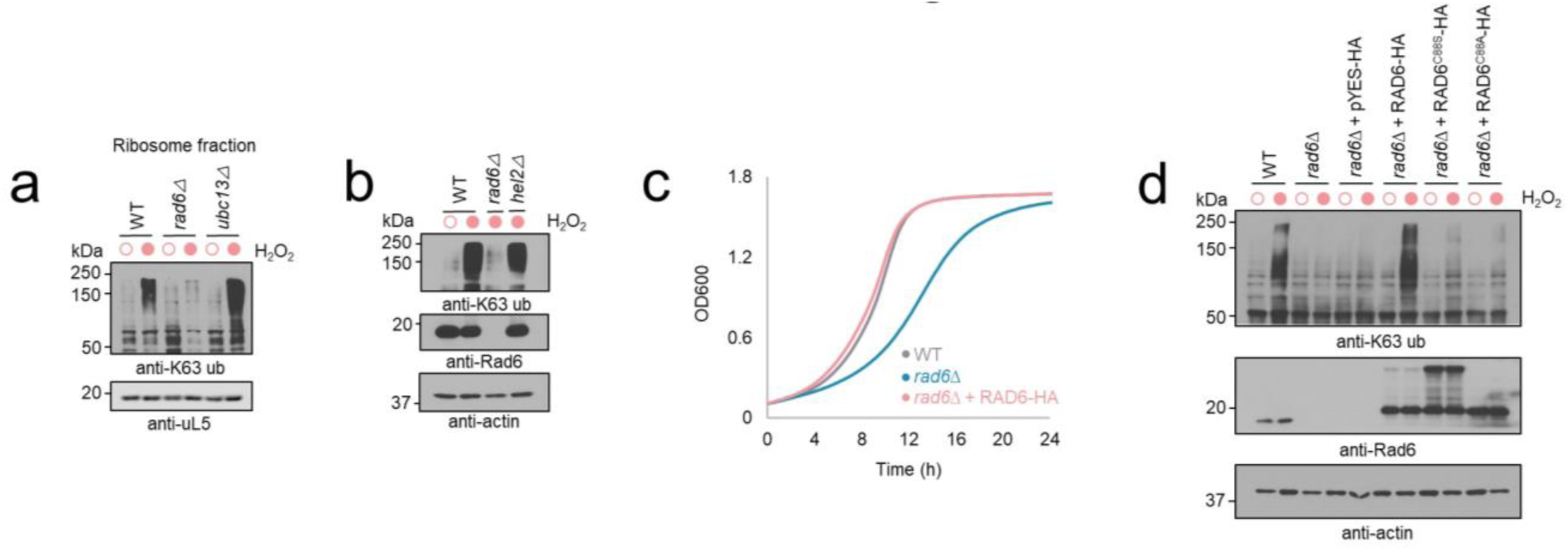
**a,b,** Deletion of *UBC13* or *HEL2* does not prevent K63 ubiquitination under stress. Immunoblot anti-K63 ubiquitin of isolated ribosomes from *rad6Δ* and *ubc13Δ* strains (**a**) and of total lysate from *rad6Δ* and *hel2Δ* (**b**) after 30 min incubation with 0.6 mM H_2_O_2_. uL5 and actin were used as loading control for ribosome isolates and total cell lysate, respectively. **c**, Expression of Rad6-HA reverts *rad6Δ* growth defect. **d,** Expression of wild type Rad6-HA supports K63 ubiquitin accumulation under stress. Mutations on Rad6 catalytic cysteine (C88) prevents K63 ubiquitin accumulation under stress. pYES corresponds to the empty vector. Anti-actin was used as loading control.

**Supplementary Figure 2.**
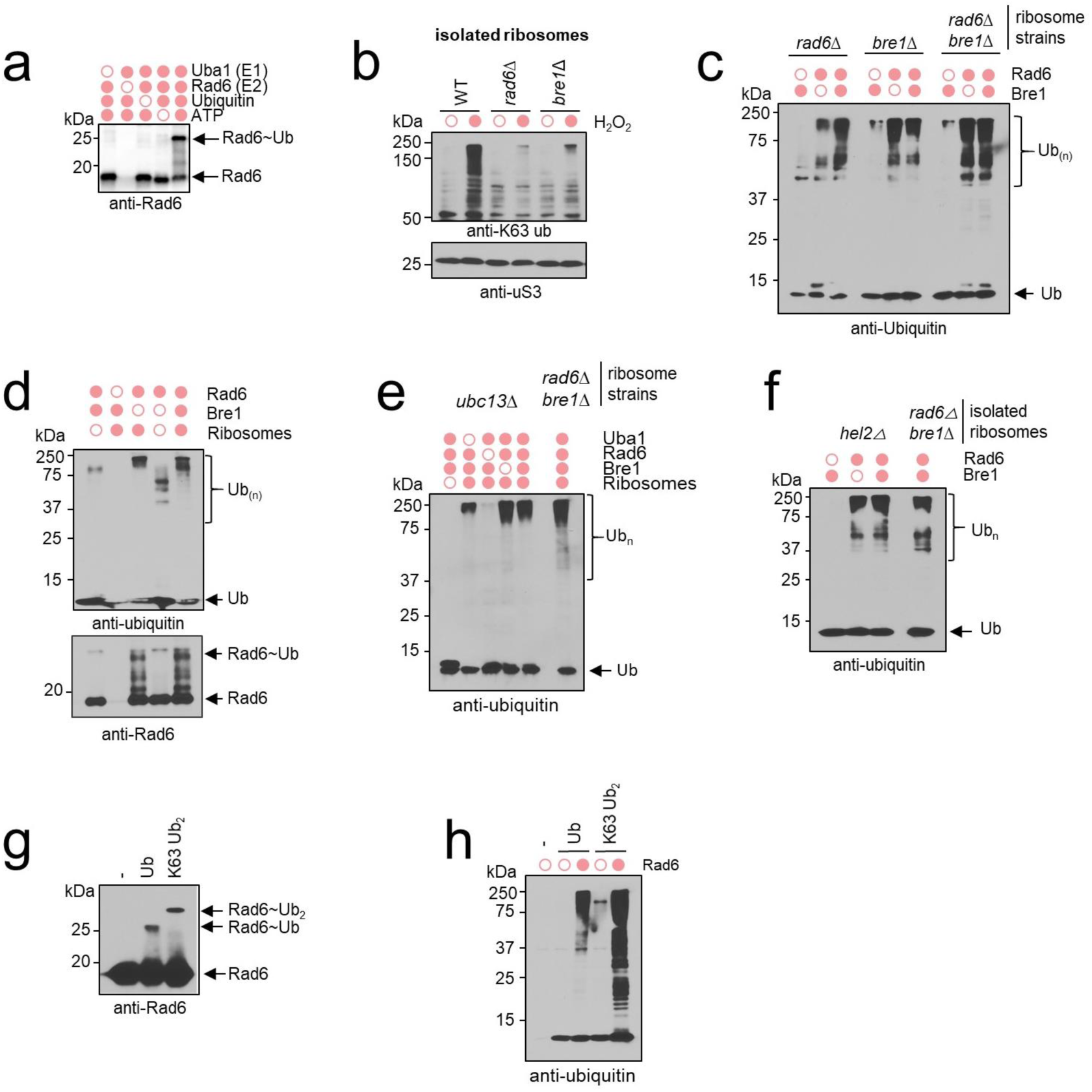
**a**, Immunoblot of Rad6 charging assay. Purified Rad6 (500 nM) was incubated with E1 (21 nM), ubiquitin (1.25 µM), and ATP (10 mM) for 30minutes at 30 °C. **b**, Immunoblot anti-K63 ubiquitin of isolated ribosomes shows that accumulation of K63 ubiquitin depends on *RAD6* and *BRE1.* Cells were treated in the presence or absence of 0.6 mM H_2_O_2_ for 30 min. Anti-uS3 was used as ribosomal loading control. **c**, Immunoblot of *in vitro* ubiquitination assay using ribosomes extracted from *rad6Δ, bre1Δ,* and the double mutant *rad6Δ bre1Δ* cells. **d**, Immunoblot of *in vitro* ubiquitination assay showing that high molecular ubiquitin conjugates are not ubiquitinated forms of Rad6, Bre1, or E1. **e,f**, Immunoblot of *in vitro* ubiquitination assay using ribosomes extracted from *ubc13Δ* and *rad6Δ bre1Δ* (**e**), *or hel2Δ* strains (**f**) to rule out Ubc13 and Hel2 contamination. **g**, Immunoblot of *in vitro* charging reaction shows Rad6 charged with K63-linked diubiquitin molecules. All lanes contain Rad6, E1, and ATP. Samples were incubated in the absence of ubiquitin, or in the presence of ubiquitin monomer (ub), or K63-linked diubiquitin (K63 Ub_2_). **h,** Immunoblot of *in vitro* ubiquitination assay of ribosomes with ub or K63 Ub_2_. All lanes contain Bre1, E1, ATP, energy generation system (creatine phosphate and creatine kinase) and ribosomes from *rad6Δ bre1Δ* strain.

**Supplementary Figure 3.**
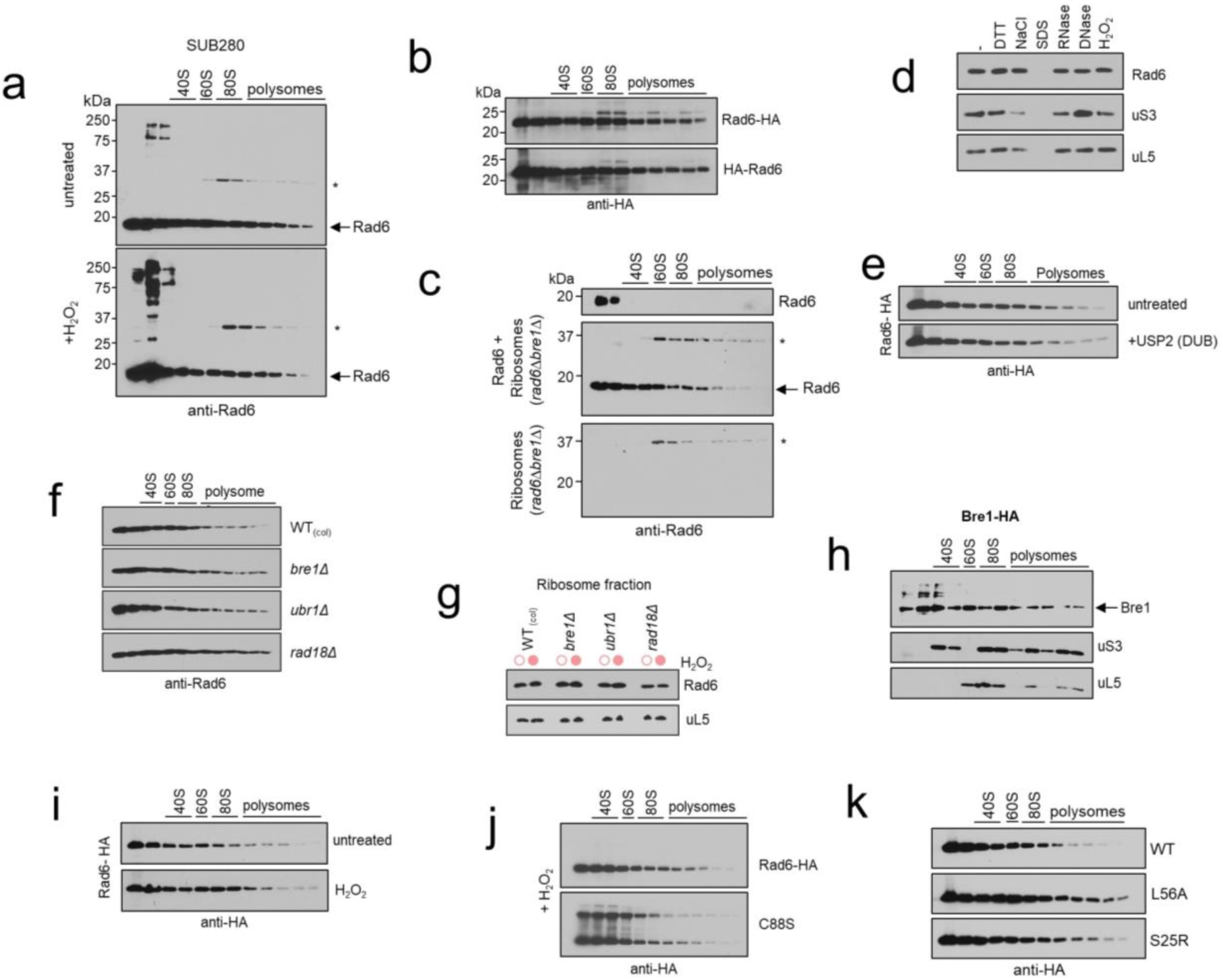
**a**, Immunoblot from sucrose sedimentation profile of wild type (WT, SUB280) cells in the presence or absence of 0.6 mM H_2_O_2_ for 30 min. **b**, anti-HA immunoblot shows that Rad6 presence in the polysome fraction of a sucrose sedimentation profile is independent of the C- or N-terminal location of the HA epitope. **c**, Immunoblot shows that purified Rad6 localizes to the polysomal fraction after incubation with ribosomes isolated from *rad6Δ* strain followed by sucrose sedimentation profile. *unspecific bands. **d**, Rad6 presence on the ribosome pellet is independent of cellular lysate incubation with 10 mM DTT, 1M NaCl, 1 mM H_2_O_2_, and DNAse (0.1 U/µg protein) and RNAse I (100 U/µg protein) treatment. 1% SDS was used to disrupt protein interactions and uS3 and uL5 were used as markers for the 40S and 60S ribosome subunits, respectively. **e,** Yeast lysate was treated with the general deubiquitin enzyme USP2 and 5 mM DTT for 30 min showing that Rad6 is not bound to ribosomes via ubiquitin. **f,g,** Rad6 presence in the polysome fraction (**f**) and ribosomal pellet (**g**) is independent of Rad6 known E3 ligase partners (Bre1, Ubr1, and Rad18)**. h**, The E3 Bre1 localizes to the polysome fraction visualized by anti-HA (Bre1) immunoblot from a sucrose sedimentation profile. uS3 and uL5 were used as markers for the 40S and 60S ribosome subunits, respectively. **i-k,** Rad6 recruitment to polysome fraction is independent of oxidative stress induced by 0.6 mM H_2_O_2_ for 30 min (**i**), Rad6 catalytic cysteine Cys88 (**j**), or Rad6 backside ubiquitin binding sites Leu56 and Ser25 (**k**).

**Supplementary Figure 4.**
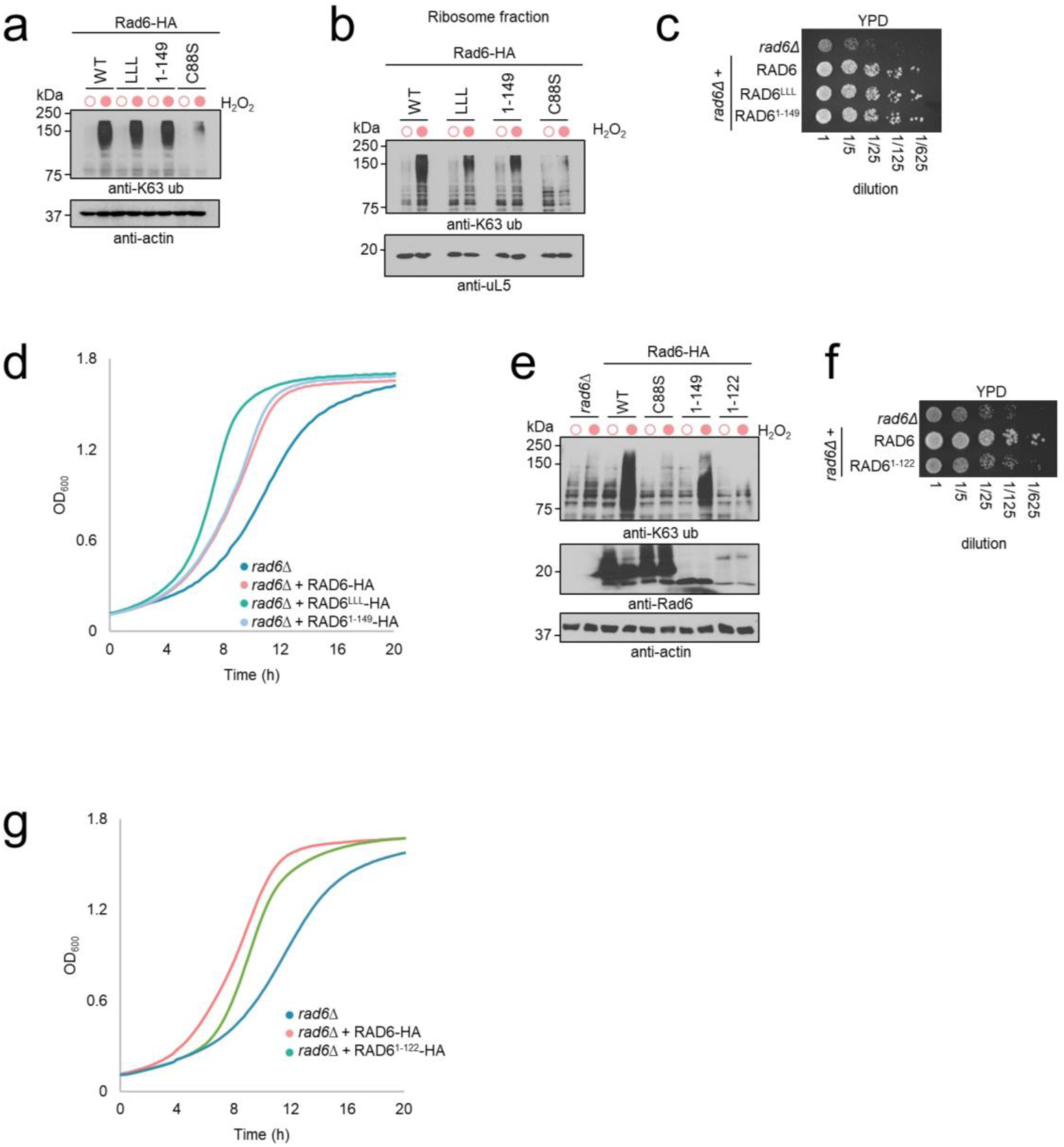
**a**,**b** Yeast cells expressing Rad6^1-149^ and Rad6^LLL^ variants are able to accumulate K63 ubiquitin in the cell lysate (**a**) and in the ribosome pellet (**b**) in response to H_2_O_2_ treatment. The mutant Rad6^C88S^ was used as negative control. Anti-actin and anti-uL5 were used as loading control. **c,d**, Expression of Rad6^1-149^ or Rad6^LLL^ reverse *rad6Δ* growth defect. **e,** Immunoblot of cell lysate from *rad6Δ* cells expressing HA-tagged wild type (WT) Rad6, Rad6^C88S^, Rad6^1-149^, or Rad6^1-122^. **f,** Expression of Rad6^1-122^ does not reverse *rad6Δ* growth defect.

**Supplementary Figure 5.**
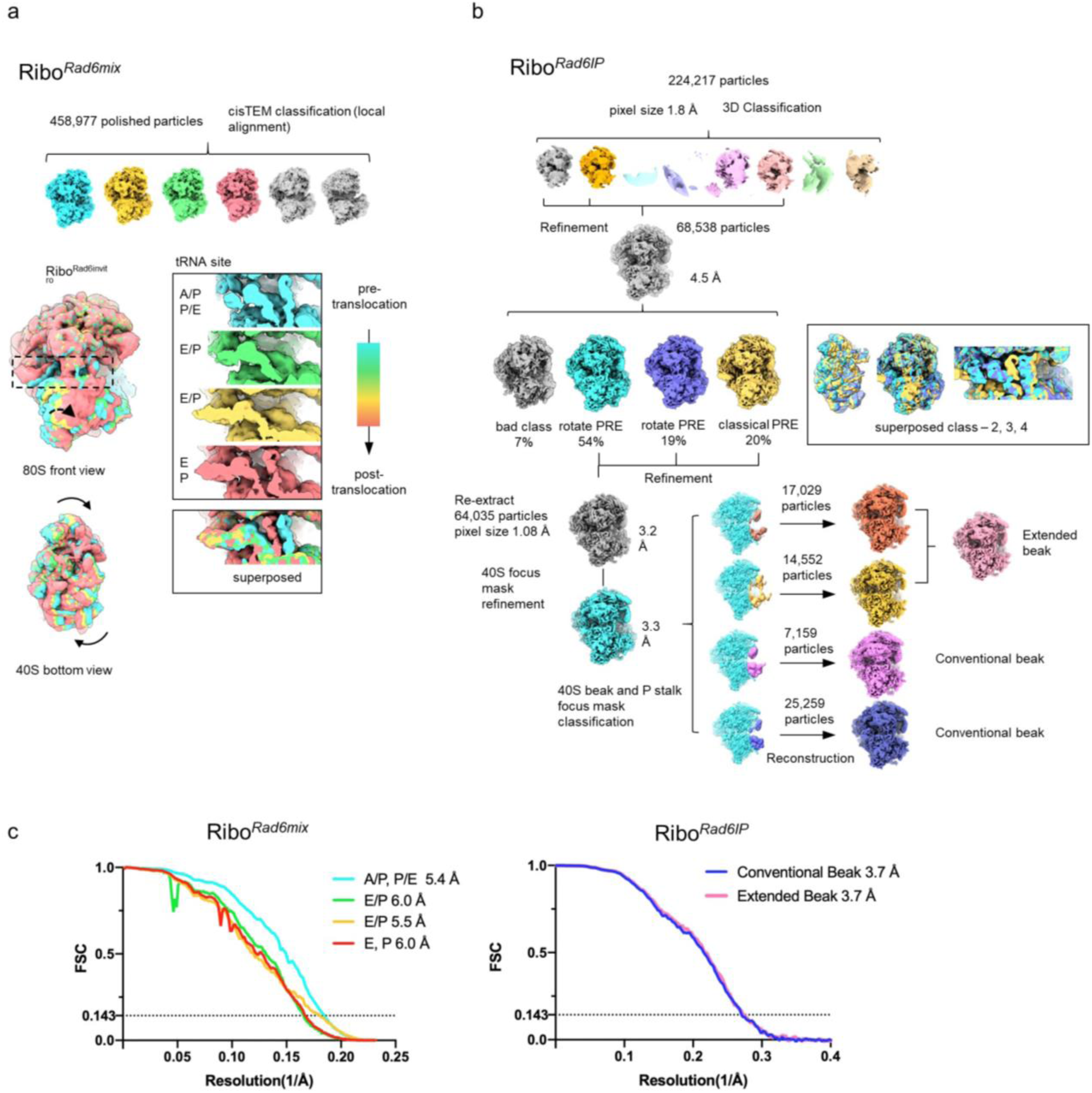
**a**, 3D classification analysis of ribosomes isolated from *rad6Δ* cells incubated *in vitro* with purified Rad6 (Ribo^Rad6mix^). Superimposed maps aligned based on the 60S showing higher variation for the 40S subunit (front view and bottom view show the rotational movement of 40S for different classes). Close up view of the decoding center shows tRNA densities and the different translation states (pre-translocation to post-translocation), depicted from cyan to salmon. **b**, 3D classification analysis of ribosomes co-immunoprecipitated with FLAG-tagged Rad6 (Ribo^Rad6IP^). Superimposed maps show the rotated pre-translocation (PRE, cyan and purple) and the classical translocation states (yellow) obtained from global classification results without alignment. The final maps resulting from the focused classification experiments are colored in pink (extended 40S beak), magenta (bad class) and blue (conventional 40S beak). **c**, Assessment of overall map resolution according to the Fourier shell correlation (FSC) 0.143-cutoff criteria for Ribo^Rad6IP^ classification results (left) and for Ribo^Rad6mix^ (right) with the conventional and extended 40S beak structures shown in Figure 2C.

**Supplementary Figure 6.**
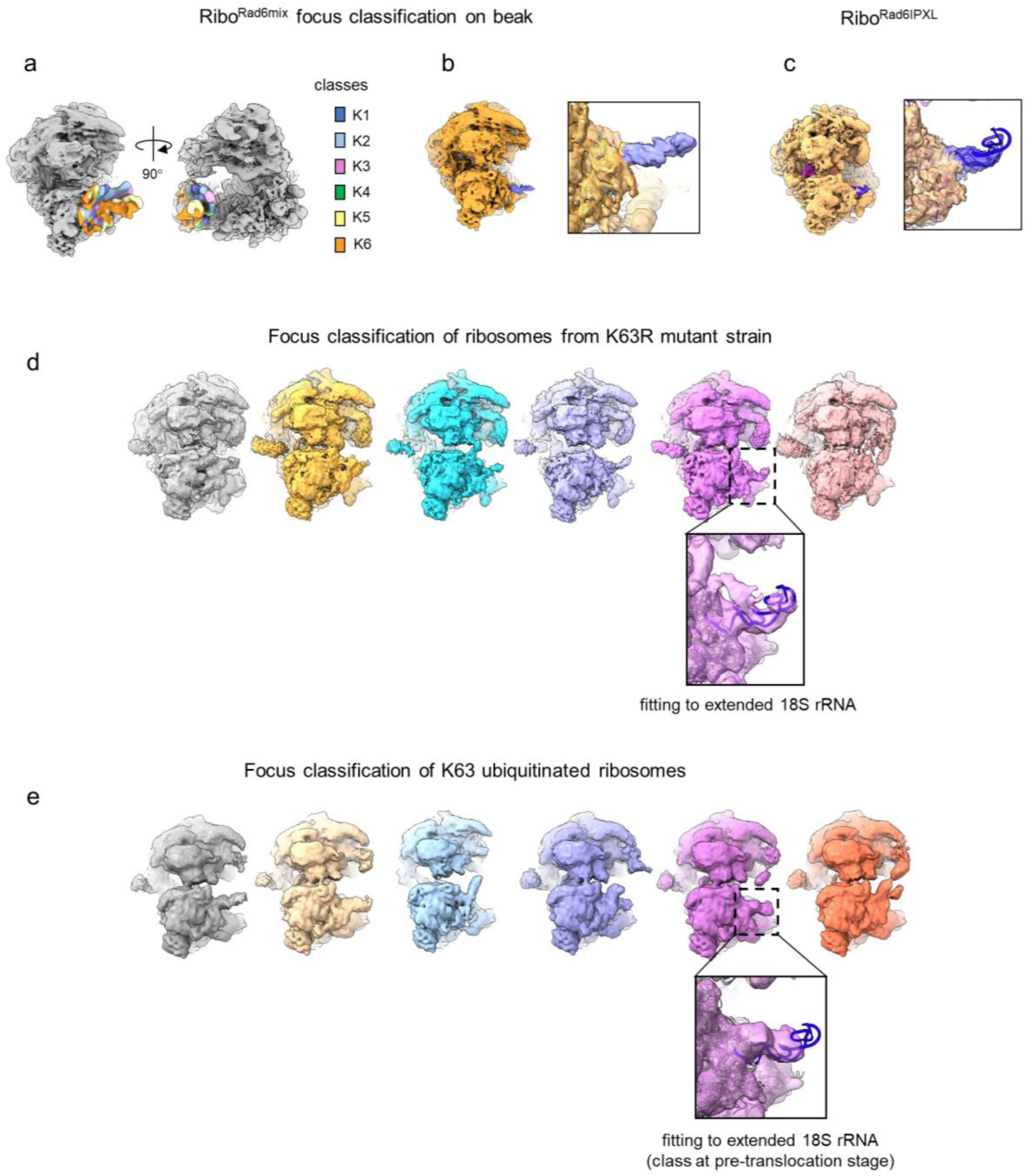
**a**, Focused classification data from Ribo^Rad6mix^ are superimposed on the 40S masked consensus of refined ribosomes. **b**, 3D reconstruction of the Ribo^Rad6mix^ K6 class, with extended 18S rRNA colored in blue. Inset shows the 40S beak density. **c**, 3D reconstruction of Ribo^Rad6IP-XL^. Inset shows the 40S beak density with extended 18S rRNA colored in blue. **d**, 3D reconstruction results of focused classification of ribosomes from the K63R ubiquitin mutant strain^1^. The back part of ribosomes is dimmed to highlight the 40S head. Zoomed-in view of the class with tRNAs reveals the extended 18s rRNA fitted into the density. **e**, Corresponding 3D reconstruction results of K63 ubiquitinated ribosomes^1^ after focused classification.

**Supplementary Figure 7.**
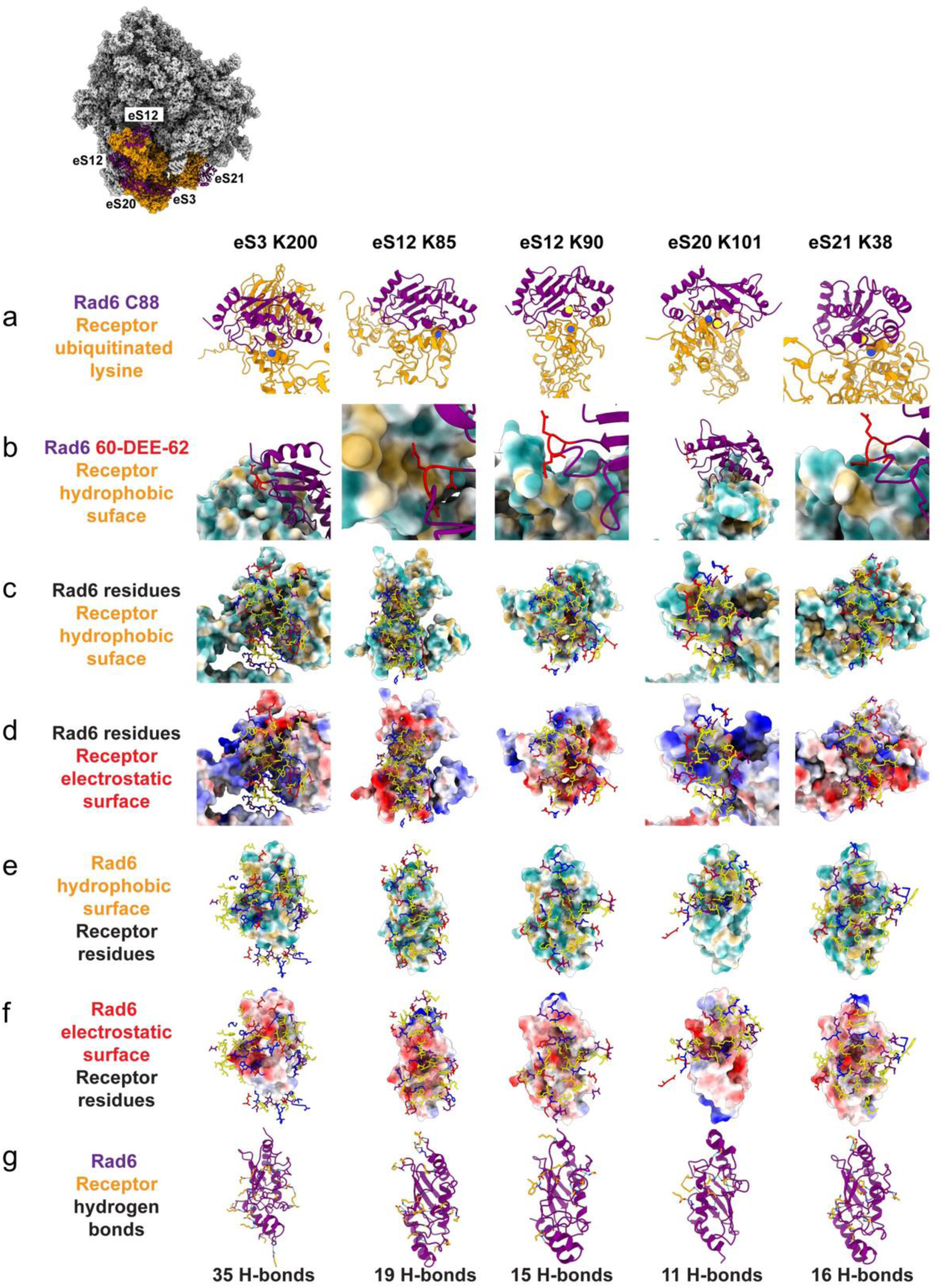
**a** Comparison of all docking predictions that are consistent with biochemical data. Rad6 was docked to ribosome receptor sites at respective ubiquitinated lysine residues: eS3 (K200), eS12 (K85), eS12 (K90), eS20 (K101), and eS21 (K38). **a,** In all cases, one of the top ten predictions from ClusPro 2.0 positioned Rad6 (purple) C88 close to the putative ubiquitinated lysine residue on the ribosome receptor (orange), suggesting a catalytically competent configuration. **b**, Placement of Rad6 60-DEE-62 (red) on the surface of the ribosome receptor colored by hydrophobicity (tan, hydrophobic; cyan, hydrophilic). Docking calculations for eS3, eS12 K85, eS12 K90, and eS21 place Rad6 60-DEE-62 close to hydrophobic patches, which agrees with the experimental result that Rad6 mutant DEE60-62LLL binds better than WT Rad6. **c**, Images show Rad6 residues (negatively charged, red; positively charged, blue; hydrophobic, yellow; and polar hydrophilic, purple) within 8 Å of the ribosome surface. **d**, Images use same color coding as above, with the ribosome surface now colored by electrostatic potential (positively charged, blue; negatively charged, red). **e**, Images display Rad6 as a surface colored by hydrophobicity, and ribosome residues within 8 Å of Rad6 shown as sticks and colored as above. **f**, Images use the same color coding as above, with Rad6 surface now colored by electrostatic potential. **g**, Images show the hydrogen bonds made between Rad6 (purple) and the ribosome surface (orange sticks). eS3 forms more hydrogen bonds than the rest of the proteins, in part due to substantially more contacts between Rad6 124-150 and the ribosome surface. In general, all docking predictions form interactions between Rad6 124-150 and the ribosome surface, which agrees with the experimental result on the importance of Rad6’s last alpha helix.

**Supplementary Figure 8.**
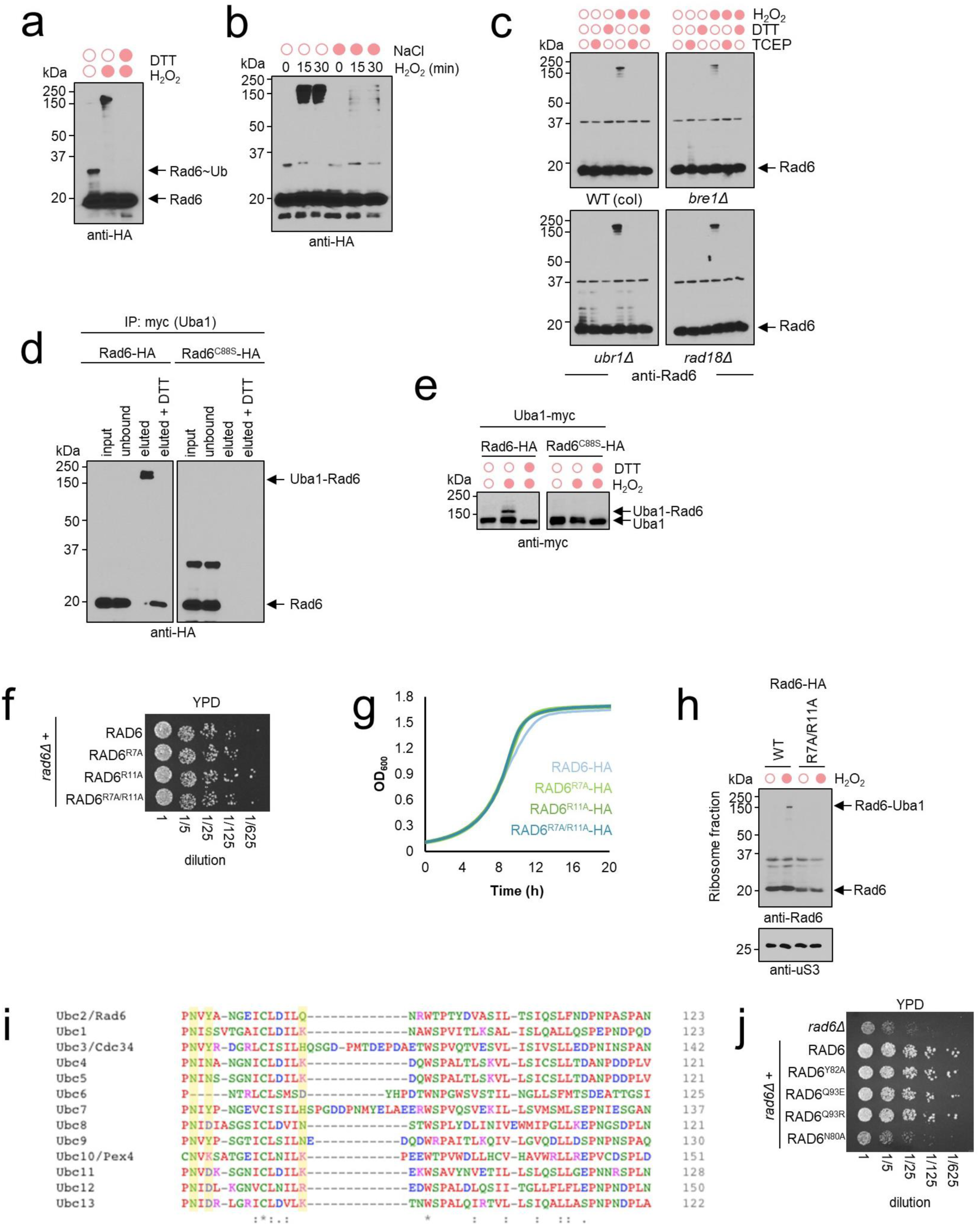
**a,** Rad6 forms heterodisulfide complex following *in vitro* incubation with 2 mM H_2_O_2_. Disulfides are reversed by 15 mM DTT. **b,** Formation of Rad6 disulfides is impacted by disruption of electrostatic interactions induced by 1 M NaCl prior to 2 mM H_2_O_2_ treatment for the designated times. **c**, Immunoblot anti-Rad6 of *in vitro* oxidation assays conducted in lysates from yeast cells deleted for *UBRE1, RAD18*, or *UBR1*. Samples were oxidized for 30 min with 2 mM H_2_O_2_ and reduced with 25 mM TCEP or DTT for 20min. (WTcol, S288c) **d,** Uba1-myc immunoprecipitation (IP) recovers complex with Rad6-HA but not with Rad6^C88S^, which is reduced by 15 mM DTT. **e**, Immunoblot anti-myc (Uba1) shows disulfide formation for Rad6 but not Rad6^C88S^, which is reduced by 15 mM DTT. **f**,**g**, Expression of Rad6^R7A^, Rad6^R11A^, and Rad6^R7A/R11A^ does not impact cell growth. **h**, Immunoblot shows that Rad6^R7A/R11A^ is present in the ribosome pellet. **i**, Sequence alignment of yeast E2s, highlighting the conservation of residues close to Rad6 catalytic cysteine. **j**, Expression of mutants Rad6^Y82A^, Rad6^Q93E^, and Rad6^Q93R^ but not Rad6^N80A^ reverse *rad6Δ* growth defect.

**Supplementary Figure 9.**
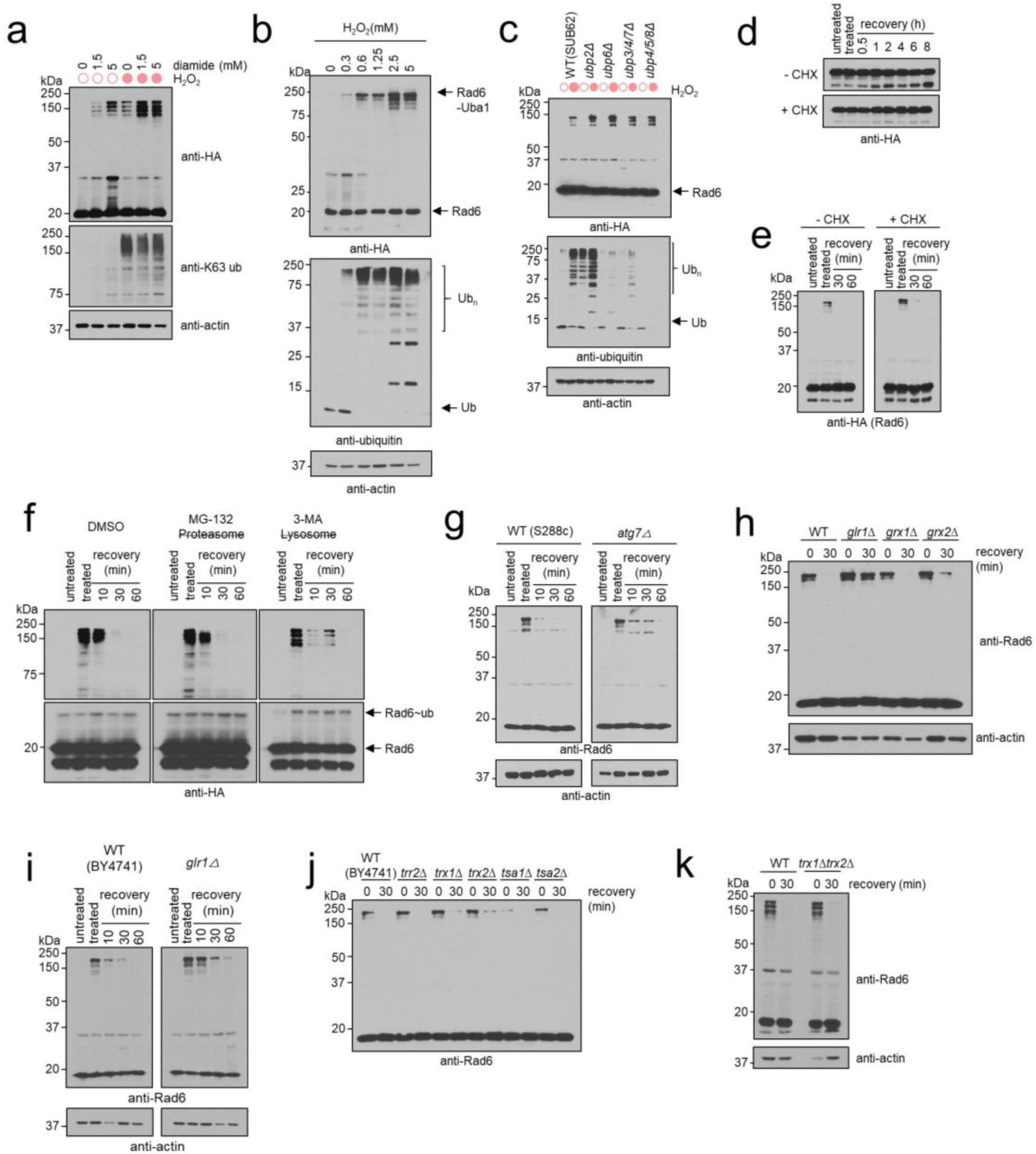
**a,** Oxidation of thiol groups by diamide for 30 min induces Rad6 disulfide formation but does not lead to increased levels of K63 ubiquitin. Samples were subsequently treated with 0.6 mM H_2_O_2_ for 30 min. **b**, Immunoblot from wild type yeast cells subjected to increased H_2_O_2_ concentrations. **c**, Deletion of DUBs that reduce the pool of free ubiquitin^2^ increase the formation of Rad6 disulfides. Cells were incubated in the presence or absence of 0.6 mM H_2_O_2_ for 30 min. **d**, Pulse-chase experiment shows that Rad6 is not degraded after stress induction. Yeast cells were pre-incubated in the presence or absence of 150 µg/ml of cycloheximide (CHX) for 30 min prior to oxidative stress induced by 0.6 mM H_2_O_2_. Cells were transferred to fresh medium containing or not CHX for up to 8 hours. Samples were DTT-reduced prior to western blot. **e,** CHX does not prevent reversal of Rad6-Uba1 disulfide. Cells were pre-incubated in the presence or absence of 150 µg/ml of cycloheximide (CHX) for 30 min prior to oxidative stress induced by 0.6 mM H_2_O_2_. Cells were transferred to to H_2_O_2_-free medium to recover in the presence or absence of CHX for up to 1 hour. **f**, Reversal of the Rad6-Uba1 complex is independent of proteasomal or lysosomal degradation. Yeast cells were grown into MPD medium^3^ and pre-incubated for 1h with the inhibitors (75 µM MG-132 of 5 mM 3-MA) prior to stress induction with 0.6 mM H_2_O_2_ for 30 min. Cells were transferred to containing the respective inhibitors. **g**, Reversal of the Rad6-Uba1 complex is independent of autophagy demonstrated by the use of the *atg7Δ* strain, which is defective in autophagosome formation ^4^. **h,i,** Immunoblot shows that deletion of glutathione reductase *(GLR1)* slows down the reduction of the Rad6-Uba1 complex. **j-k,** Immunoblot shows that reduction of the Rad6-Uba1 complex is not impacted by the deletion of genes of the thioredoxin family. Anti-actin was used as loading control.

**Supplementary Figure 10.**
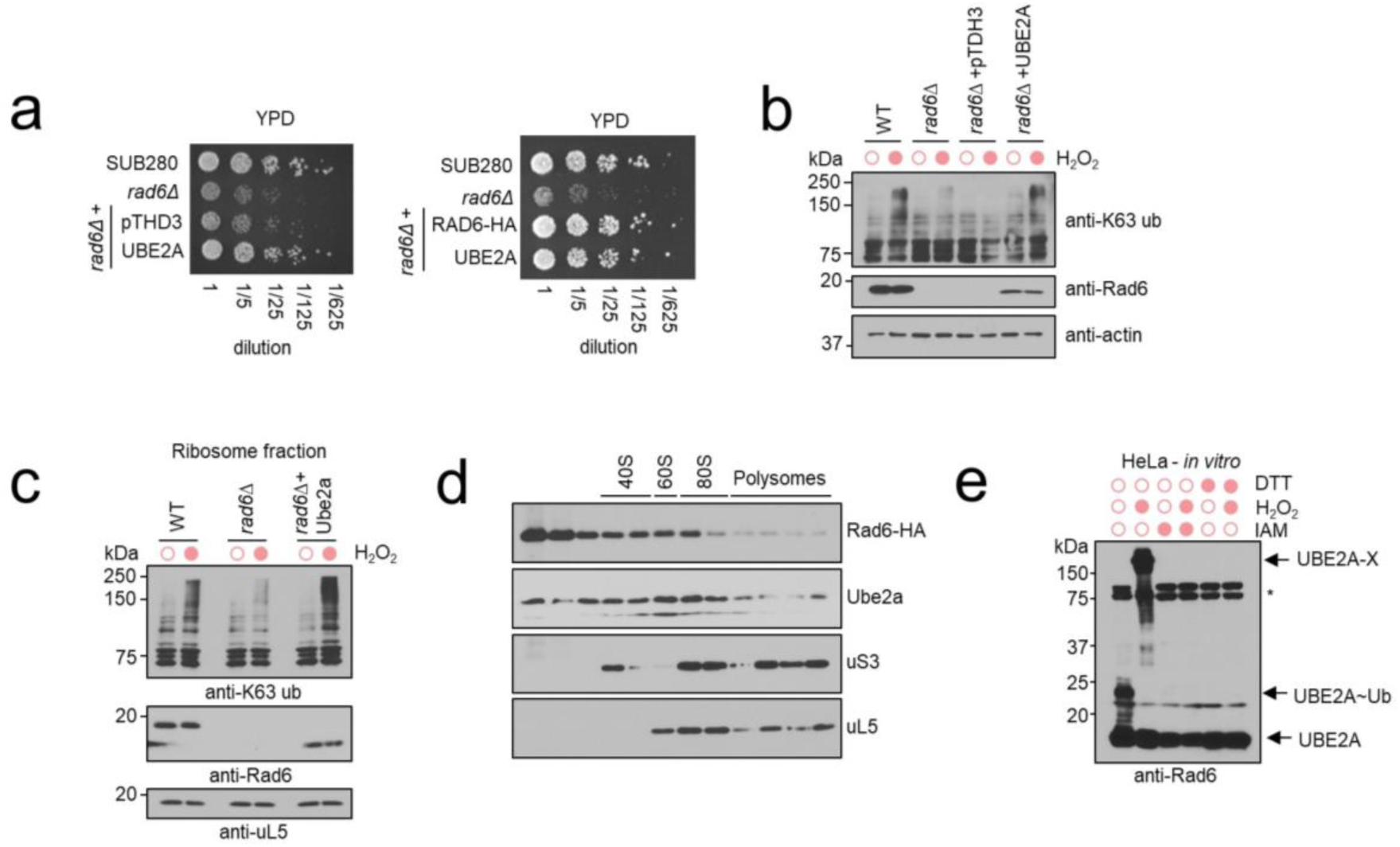
**a**, Expression of UBE2A in yeast cells reverts *rad6Δ* growth defect. **b,** Expression of UBE2A supports *rad6Δ* K63 ubiquitin accumulation under 0.6 mM H_2_O_2_ by immunoblotting cell lysate. **c,** Immunoblotting showing that UBE2A expressed in yeast cells localizes to the ribosome fraction. pTDH3 represents the empty vector. Anti-actin and anti uL5 were used as loading control. **d**, anti-Rad6 immunoblot shows UBE2A presence in the polysome fraction of a sucrose sedimentation profile. **e**, HeLa cell lysate incubated *in vitro* with 20 mM cysteine alkylator iodoacetamide (IAM), oxidized with 2 mM H_2_O_2_, or reduced with 15 mM dithiothreitol (DTT). anti-Rad6 (UBE2A) immunoblot shows the formation of reversible disulfides. * unspecific bands.

**Supplementary Figure 11.**
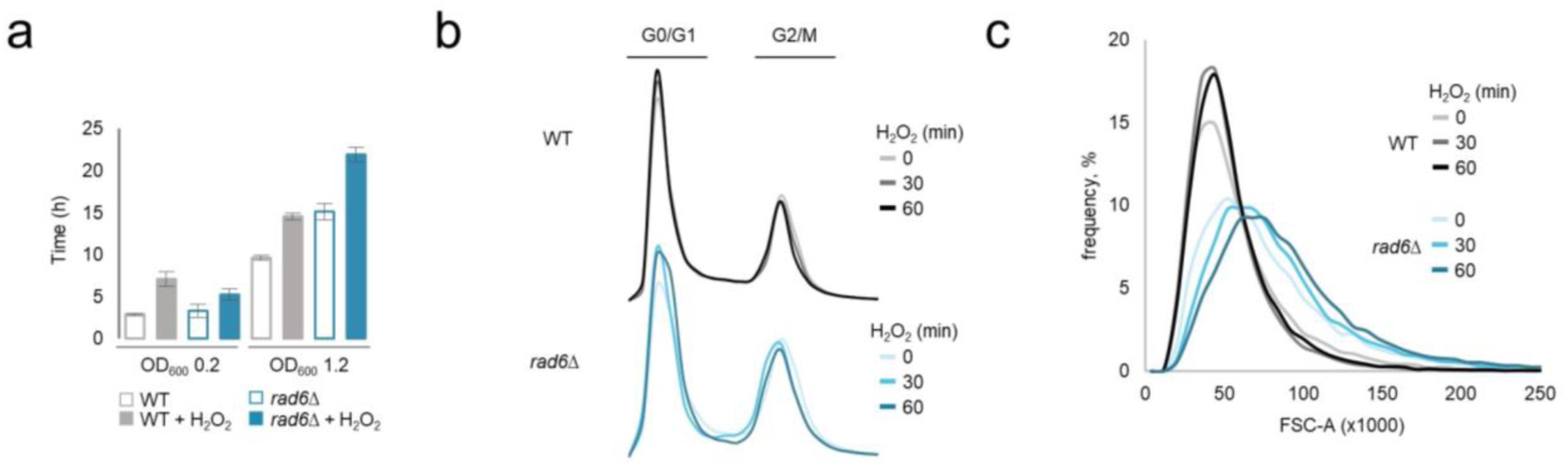
**a,** Average time required for yeast cells from wild type (WT) and *rad6Δ* strains to reach OD_600_ of 0.2 and 1.2 starting from OD_600_ of 0.05 in the presence or absence of H_2_O_2_ stress. Data was determined from three biological replicates. A complete growth curve is presented in Figure 6C. **b,** Cell cycle profile from flow cytometric analyses of DNA contents and **c**, forward scatter (FSC) plots of WT and *rad6Δ* yeast strain under 0.6 mM H_2_O_2_ for 30 and 60 min as described in Materials and Methods.

